# Th17 immunity in the colon is controlled by two novel subsets of colon-specific mononuclear phagocytes

**DOI:** 10.1101/2021.01.28.428597

**Authors:** Hsin-I Huang, Mark L. Jewell, Nourhan Youssef, Min-Nung Huang, Elizabeth R. Hauser, Brian E. Fee, Nathan P. Rudemiller, Jamie R. Privratsky, Junyi J. Zhang, Estefany Reyes, Donghai Wang, Gregory A. Taylor, Michael D. Gunn, Dennis C. Ko, Donald N. Cook, Vidyalakshmi Chandramohan, Steven D. Crowley, Gianna Elena Hammer

## Abstract

Intestinal immunity is coordinated by specialized mononuclear phagocyte populations, constituted by a diversity of cell subsets. Although the cell subsets constituting the mononuclear phagocyte network are thought to be similar in both small and large intestine, these organs have distinct anatomy, microbial composition, and immunological demands. Whether these distinctions demand organ-specific mononuclear phagocyte populations with dedicated organ-specific roles in immunity are unknown. Here we implement a new strategy to subset murine intestinal mononuclear phagocytes and identify two novel subsets which are colon-specific: a macrophage subset and a Th17-inducing dendritic cell (DC) subset. Colon-specific DCs and macrophages co-expressed CD24 and CD14, and surprisingly, both were dependent on the transcription factor IRF4. Novel IRF4-dependent CD14^+^CD24^+^ macrophages were markedly distinct from conventional macrophages and failed to express classical markers including CX3CR1, CD64 and CD88, and surprisingly expressed little IL-10, which was otherwise robustly expressed by all other intestinal macrophages. We further found that colon-specific CD14^+^CD24^+^ mononuclear phagocytes were essential for Th17 immunity in the colon, and provide definitive evidence that colon and small intestine have distinct antigen presenting cell requirements for Th17 immunity. Our findings reveal unappreciated organ-specific diversity of intestine-resident mononuclear phagocytes and organ-specific requirements for Th17 immunity.

## Introduction

The intestinal immune system must manage a diversity of antigens such as food, intestine-resident microbiota, and intestinal pathogens. Immunity to these is coordinated by specialized populations of mononuclear phagocytes, including dendritic cells (DCs) and macrophages. These two subsets have distinct functions, with macrophages primarily serving as phagocytic scavengers and robust producers of IL-10, and DCs primarily serving as antigen presenting cells (APC) for naïve T cells^1–4^. An additional layer of functional specialization is embedded within intestinal DC and macrophage populations since both cell types are composed of discrete subsets, some of which are specific to the intestine and do not exist in other tissues. Evidence suggests that each DC and macrophage subset is functionally distinct and therefore, accurate and complete identification of these subsets is a major priority, since these mononuclear phagocytes are key for pathogen defense, intestinal homeostasis, and also, the aberrant activity of intestinal DCs and macrophages is linked to pathogenesis of inflammatory bowel disease and colorectal cancer^5–10^.

Accurate identification of mononuclear phagocyte subsets is fundamental to understanding intestinal immunity and based on several landmark studies^10–15^, murine macrophages and DCs (within MHC-II^+^CD11b^+^/CD11c^+^ populations) are commonly distinguished on the basis of CD64 or CD14. Both markers are highly expressed on intestinal macrophages, whereas DCs must meet two criteria: 1) Negative for CD14 and CD64, and 2) Positive for CD24 or CD26^10–15^. Although these subsetting strategies are widely used, previous studies validating these approaches largely focus on small intestine and it remains to be thoroughly investigated whether these strategies are equally as effective in the colon^10–15^.

Murine intestinal macrophages exist in at least three subsets based on expression of CD4 and Tim-4, with Tim-4^+^ macrophage subsets having the slowest turnover rate^16^. Single-cell gene expression suggests additional diversification exists among these macrophage subsets, some of which is driven in a microbiota-dependent fashion^17^. While each macrophage subset has a unique transcriptome, all are thought to produce robust amounts of IL-10^16^. Intestinal DCs are among the most diverse in the body and there are three subsets of conventional DCs (cDCs) currently identified: CD103^+^CD11b^−^, CD103^+^CD11b^+^, and CD103^−^CD11b^+^ DCs^3,15,18,19^. Consistent with defining criterion for the cDC lineage, all three subsets express the cDC-specific transcription factor *Zbtb46*, are FLT3L-depdendent, and develop from pre-DC precursors^20^. cDCs instruct CD4 T cells to respond to intestinal microbes and these APC functions are highly subset specific, particularly so for DCs that instruct IL-17-producing CD4 T cells (Th17)^6,21–24^. Th17-inducing DC functions are exclusive to IRF4^+^ intestinal DC subsets (currently identified to be CD103^+^CD11b^+^ and CD103^−^CD11b^+^ cDC subsets), and the absence of IRF4-expressing DCs from small intestine results in concomitant loss of Th17 cells from this organ^10,14,25,26^.

Since all DC and macrophage subsets currently identified can be found in both small and large intestine, it is commonly thought that these organs have similar APC requirements for Th17 immunity, although this has not been formally tested. Small and large intestine have several remarkable differences, including structural distinctions of the epithelial villi and mucus layer^27–29^ and immunological distinctions such as the predominant localization of Paneth cells to small intestine^30^, and the absence of Peyer’s patches in the colon^31^. Moreover, microbial abundance and composition are distinct, with the total microbial burden and the abundance of anaerobic microbes being significantly higher in colon as compared to small intestine^32,33^. While these distinctions highlight the unique immunological demands of each intestinal organ, whether these demands necessitate organ-specific mononuclear phagocyte populations with dedicated organ-specific roles in immunity is unknown. Here we identify two novel populations of mononuclear phagocytes, a cDC subset and an atypical IL-10^low^ macrophage subset, which are colon-specific and do not reside in small intestine. We further show that these novel colon-specific APCs are IRF4-dependent and essential for Th17 immunity in the colon. These findings define new subsets of mononuclear phagocytes and surprisingly reveal the existence of organ-specific APCs for immune surveillance in the intestine.

## Results

### Subsetting by CD14, CD24, and CD88 identify a novel population of colon-specific APCs

Among intestinal mononuclear phagocytes the largest amount of heterogeneity exists among MHC-II^+^ populations that express either CD11c or CD11b (referred to here as MHC-II^+^CD11c/CD11b^+^ APCs).

These populations are a mixture of macrophages, monocytes differentiating into macrophages, and DCs. To distinguish these cell types in intestinal lamina propria we gated on MHC-II^+^CD11c/CD11b^+^ APCs and used CD14 and CD24, standard markers commonly used to distinguish DCs (CD14^−^CD24^+^) and macrophages (CD14^+^CD24^−^) in murine intestine^10–12,14,15^ (Fig 1a,b; see Fig S1 for complete gating strategy). Our analysis of small intestine aligned with previous reports in that CD14 and CD24 expression was almost entirely mutually exclusive, with very few double positive cells (CD14^+^CD24^+^) (Fig 1c). While CD14^+^CD24^+^ APCs were few in small intestine, colon surprisingly contained a robust CD14^+^CD24^+^ population (Fig 1d). Intestinal CD14^+^CD24^+^ APCs have not been previously described and because the abundance of these was markedly distinct between small intestine and colon, we set out to define the cellular lineage of these populations. We first assessed CD64, a well-established marker of intestinal macrophages^11,13,15^. In alignment with previous reports on small intestine^11,13,15^ we found all small intestinal CD14^+^ APCs co-expressed CD64 and thus, these were categorically macrophages (Fig 1e). CD64 was also expressed by the majority of CD14^+^ APCs in the colon, including some CD14^+^CD24^+^APCs, suggesting that some colon-resident cells with the CD14^+^CD24^+^ phenotype belonged to the macrophage lineage (Fig 1f). Surprisingly however, colon also contained CD14^+^CD24^+^ populations that were CD64^negative^. These CD14^+^CD24^+^CD64^negative^ cells appeared to be exclusive to the colon (Fig 1e,f).

**Figure 1.**
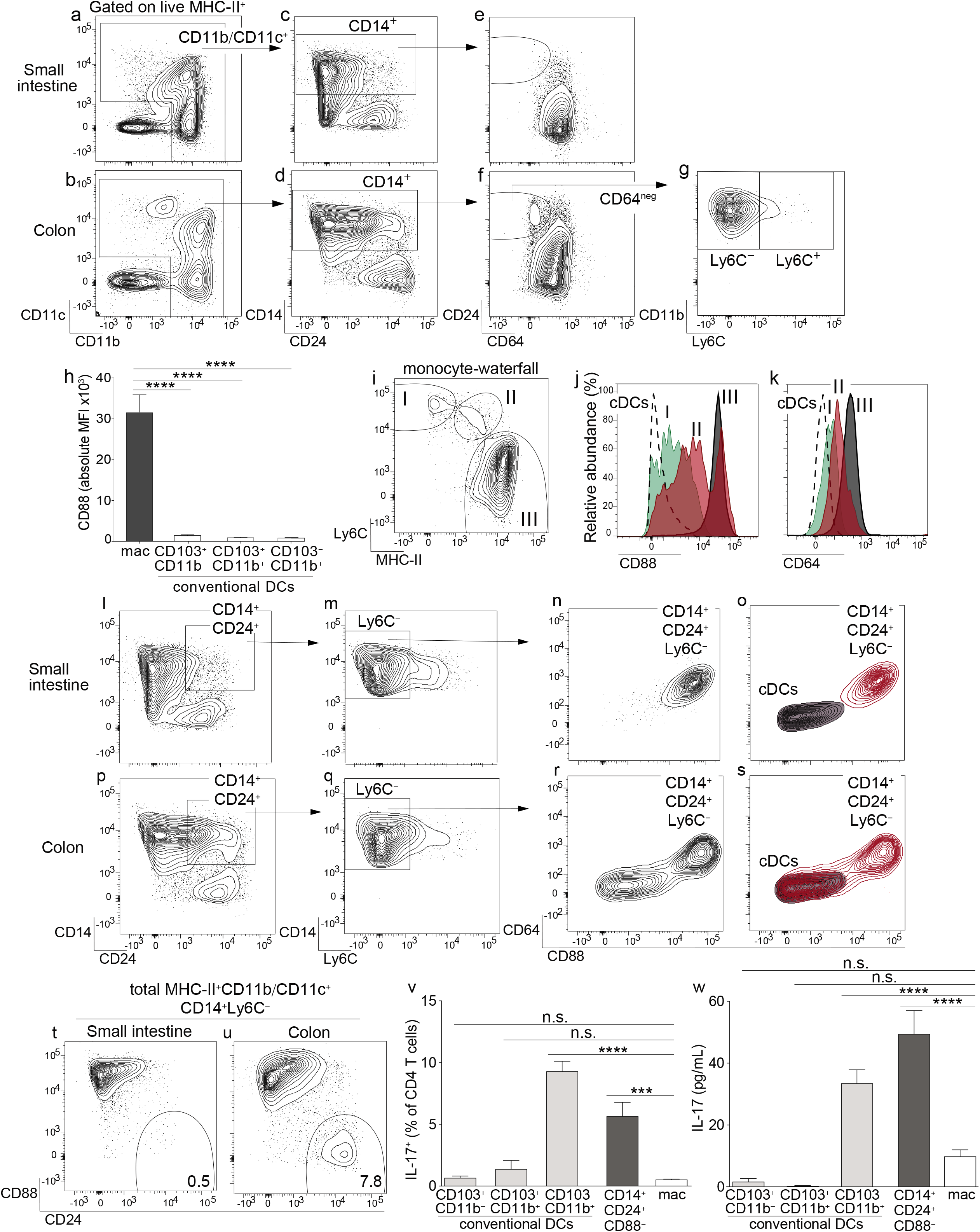
Identification of a novel population of CD14^+^CD24^+^CD88^−^ mononuclear phagocytes that reside in colon and are exclusive to this intestinal organ. Representative plot gated on live MHC-II^+^ APCs from small intestine lamina propria (SI-LP) (a) or colon lamina propria (C-LP) (b). (c-d) CD11b/CD11c^+^ APCs from SI-LP (c, gated in (a)) or C-LP (d, gated in (b)) evaluated for CD14 and CD24 expression. (e-f) CD14^+^ populations from SI-LP (e, gated in (c)) or C-LP (f, gated in (d)) evaluated for CD24 and CD64 expression. (g) The CD64^negative^ population from C-LP (gated in (f)) evaluated for Ly6C expression. (h) Mean fluorescence intensity (MFI) of surface CD88 expressed by SI-LP macrophages and each of the three cDC subsets (See Fig S1 for macrophage and cDC gating strategy). Each dot represents one mouse. (i) Representative plot gated on total CD11b^+^CD24^−^ cells from SI-LP depicting populations representing the developmental transition of monocytes into intestinal macrophages (termed the monocyte-waterfall): (I) Ly6C^+^MHC-II^−^ monocytes; (II) Ly6C^+^MHCII^+^ transitioning monocytes; (III) Ly6C^−^MHCII^+^ mature macrophages. (j-k) Gated populations shown in (i) and total cDCs were evaluated for CD88 (j) and CD64 expression (k): Dashed line, cDCs; (I) Green, monocytes; (II) Red, transitioning monocytes; (III) Gray, mature macrophages. (l) SI-LP APCs gated as in (a) were evaluated for CD14 and CD24. (m) CD14^+^CD24^+^ populations gated in (l) were evaluated for Ly6C. (n) Ly6C^−^ populations gated in (m) were evaluated for CD64 and CD88. (o) Contour plot overlay of SI-LP CD14^+^CD24^+^Ly6C^−^ populations (grey, gated in (m)) and SI-LP cDCs (grey). (p-s) C-LP APCs were gated and analyzed as in (l-o). (t-u) Representative plots of total live MHC-II^+^CD11b/CD11c^+^CD14^+^Ly6C^−^ APCs from SI-LP (t) or C-LP (u) evaluated for CD88 and CD24 expression (See Fig S3 for gating strategy). (v,w) C-LP mononuclear phagocytes of the indicated subset were co-cultured 1:1 with naïve OT-II CD4 T cells for 4 days. (v) T cells from each co-culture were restimulated and evaluated for the percentage of IL-17^+^ OT-II T cells. (w) The abundance of IL-17 protein in co-culture supernatant prior to stimulation was analyzed in parallel. Data in (v,w) is combined from three independent experiments each containing sorted APCs from at least 5 mice. Error bar represents mean ± SEM. Data in (a-u) is representative of >40 individual mice. \**p* < 0.05, \*\*\*\**p* < 0.0001, n.s., not significant (one-way ANOVA with Tukey’s post hoc test).

We considered the possibility that CD14^+^CD24^+^CD64^negative^ APCs were transitioning monocytes undergoing the “monocyte-waterfall”, the process wherein Ly6C^+^ monocytes develop into MHC-II^+^ intestinal macrophages^2,13,34^. In contrast to this hypothesis, very few CD14^+^CD24^+^CD64^negative^ cells expressed Ly6C, suggesting that these were not transitioning monocytes (Fig 1g). Identification of transitioning monocytes based on their co-expression of MHC-II and Ly6C is standard in the field, but a potential limitation of this approach lies in the fact that Ly6C is downregulated during the monocyte-waterfall^13,35^. We thus sought to examine markers that were upregulated during the monocyte-waterfall and invariably expressed at high levels on mature intestinal macrophages. We reasoned that this approach would capture still-developing monocytes/macrophages that had downregulated Ly6C to below the limit of detection. We further reasoned that this approach would capture all macrophage subsets, including those that do not have monocytic origins, although these are expected to be a very minor population in the intestine of adult mice^34^. To this end we tested the utility of CD88, a molecule recently shown to be highly expressed on intestinal macrophages but absent on intestinal cDCs^36^. We confirmed the findings of that report and additionally found that all mature intestinal macrophages, including CD4^+^ and Tim-4^+^ subsets, expressed uniformly high levels of CD88 (Fig 1h, Fig S2). Importantly, CD88 was highly upregulated during the monocyte-waterfall, supporting the utility of CD88 to distinguish both mature macrophages and transitioning monocytes committed to the macrophage lineage (Fig 1i,j). Upregulation of CD88 during the monocyte-waterfall, and the absence of CD88 on intestinal cDCs, very closely mirrored the expression pattern of CD64, a validated marker of intestinal macrophages (Fig 1k, Fig S2)^11,13^. Although in comparison to CD64 (peak intensity of ~10^3^), CD88 staining was far more robust (peak intensity of ~10^5^) and also, it was clear that CD88 expression on most transitioning monocytes and all mature macrophages was well above that of cDCs (Fig 1h,j,k). Taken together, these results suggest CD88 was a preferable marker with which to distinguish intestinal macrophages, as well as still-developing monocytes committed to the macrophage lineage.

Based on these results we tested for CD88 expression on CD14^+^CD24^+^ APCs. Although CD14^+^CD24^+^ APCs were a very minor population in small intestine (Fig 1c), we nevertheless gated on these to elucidate whether small and large intestine truly had organ-specific differences in their composition of APC subsets, as suggested by our previous results (Fig 1e,f). We gated on CD14^+^CD24^+^Ly6C^−^ APCs from small intestine and found all of these expressed very high levels of CD88 (Fig 1l-n). These also co-expressed CD64, consistent with our previous results (Fig 1e). By virtue of CD64 and CD88 expression, all CD14^+^CD24^+^Ly6C^−^ APCs from small intestine were readily distinguishable from cDCs (Fig 1o). Taken together, these results indicate that the few CD14^+^CD24^+^ APCs found in small intestine belonged to the macrophage lineage.

In agreement with small intestine, many colon-resident CD14^+^CD24^+^Ly6C^−^ APCs co-expressed CD64 and CD88 (Fig 1p-r). However, colon also contained a striking population that was both CD88^negative^ and CD64^negative^. That neither CD88, CD64 or Ly6C was expressed by CD14^+^CD24^+^CD88^−^ APCs strongly suggests that these were neither transitioning monocytes nor conventional macrophages. In fact, if not for robust CD14 expression, colon-resident CD14^+^CD24^+^CD88^−^ APCs were indistinguishable from cDCs (Fig 1s). Comparative analysis of total CD14^+^ APCs from colon and small intestine readily distinguished CD14^+^CD24^+^CD88^−^ APCs, and further demonstrated that these were highly colon-specific (Fig 1t,u; gating strategy shown in Fig S3). Colon-resident CD14^+^CD24^+^CD88^−^ APCs expressed CD11b and CD11c, but were negative for CD103, Ly6G, CD64, F4/80, Tim-4, and CD4 (Fig S3).

Given that colon-specific CD14^+^CD24^+^CD88^−^ APCs expressed MHC-II but failed to align with all currently known intestinal cDC or macrophage subsets, we tested whether these were capable of activating naïve CD4 T cells. We sorted CD14^+^CD24^+^CD88^−^ APCs and tested their APC functions in co-cultures with naïve OT-II CD4 T cells. We also tested each of the three cDC subsets (CD103^+^CD11b^−^, CD103^+^CD11b^+^ and CD103^−^CD11b^+^) and conventional macrophages (CD14^+^CD24^−^) sorted from the same mice. In these co-cultures, instruction of IL-17-producing CD4 T cells (Th17) was robust in only two APC subsets: CD103^−^CD11b^+^ cDCs, which are known to possess Th17-inducing functions^3^, and also, the novel CD14^+^CD24^+^CD88^−^ APC subset (Fig 1v). That the CD14^+^CD24^+^CD88^−^ subset possessed Th17-inucing APC functions was further supported by the abundance of IL-17 cytokine in co-culture supernatant (Fig 1w). CD14^+^CD24^+^CD88^−^ APCs also instructed IFNγ-producing CD4 T cells (Th1), although in these experiments Th1-inducing functions were not significantly different between APC subsets (Fig S4). Taken together with our findings above, CD14^+^CD24^+^CD88^−^ APCs possess Th17-inducing functions and are phenotypically distinct from conventional macrophages and all known cDC subsets. Most strikingly, these novel Th17-inducing APCs are colon-specific and do not exist in small intestine.

### Colon-specific CD14^+^CD24^+^CD88^−^ APCs are a heterogenous population composed of two subsets: one dendritic cell, the other an atypical macrophage

We set out to establish whether CD14^+^CD24^+^CD88^−^ APCs belonged to the DC or macrophage lineage. To this end we tested for expression of CX3CR1, which is highly expressed on intestinal macrophages, but is either absent or weakly expressed on intestinal DCs^3,15,18,37^. Using CX3CR1-GFP mice we found that CD14^+^CD24^+^CD88^−^ APCs were a mixture of CX3CR1^intermediate^ and CX3CR1^negative^ cells (Fig 2a). To resolve these populations further we tested for CD26, since CD26 is expressed by all known intestinal cDCs and has also been proposed to be a universal cDC marker, expressed on cDCs from different tissues and across mouse strains^12,15,36^. Among intestinal mononuclear phagocytes we confirmed that CD26 was expressed by cDCs and was not expressed by macrophages, or by any subsets within the monocyte-waterfall (Fig S5). Interestingly, CD26 distinguished CD14^+^CD24^+^CD88^−^ APCs into two discrete populations (Fig 2b; See Fig S5 for isotype control). Only the CD26^+^ population expressed CX3CR1, as a heterogeneous mixture of CX3CR1^intermediate^ and CX3CR1^negative^ cells (Fig 2c). This heterogeneity resembled CX3CR1^int/neg^ expression by the CD103^−^CD11b^+^ cDC subset^3^ (Fig S5).

**Figure 2.**
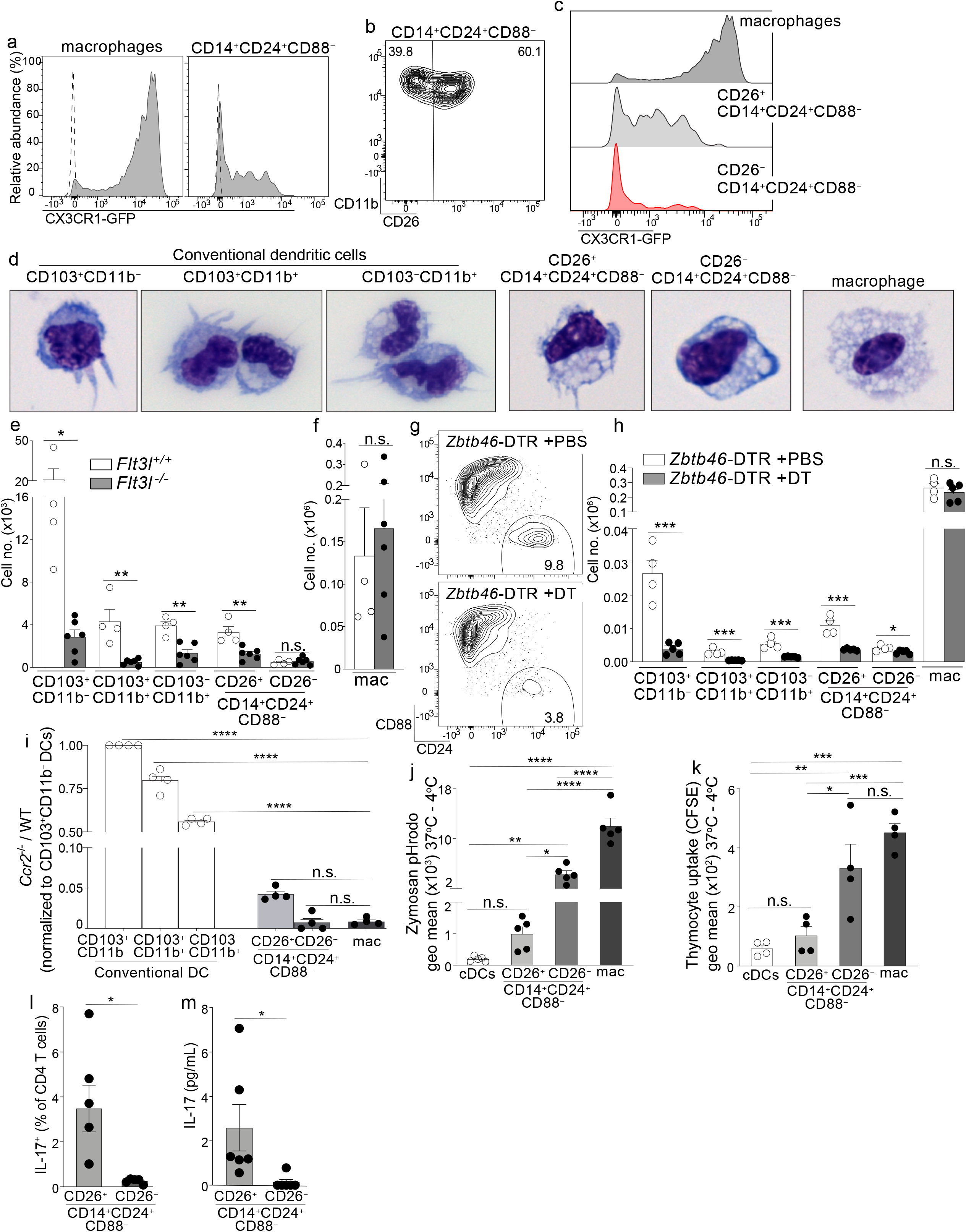
Colon-specific CD14^+^CD24^+^CD88^−^ APCs are composed of a CD26^+^CX3CR1^int/neg^ cDC subset and an atypical CD26^−^CX3CR1^−^ macrophage subset. (a) Representative histogram of CX3CR1-GFP expression by C-LP macrophages and CD14^+^CD24^+^CD88^−^ APCs (gated as in Fig 1u). Dashed line represents GFP expression by the respective populations from wild-type, GFP negative control mice. (b) Representative plot of CD26 expression on C-LP CD14^+^CD24^+^ CD88^−^ APCs. (c) Comparative CX3CR1-GFP expression by the indicated C-LP population. (d) Representative light microscopy image of sorted cells from the indicated C-LP mononuclear phagocyte subset. Data is representative of two independent experiments. (e-f) Absolute cell number of the indicated subset of C-LP mononuclear phagocyte (e), and macrophages (f) in *Flt3l*^+/+^ and *Flt3l^−/−^* mice. Data was combined from two independent experiments. (g-h) Hematopoietic chimera reconstituted with *Zbtb46*-DTR bone marrow were injected with diphtheria toxin (DT) to deplete *Zbtb46*-expressing cells. Control chimera injected with PBS alone were analyzed in tandem. Representative flow plot gated as in Fig 1u indicating the relative abundance of CD14^+^CD24^+^CD88^−^ APCs (g), or the absolute number (h) of the indicated mononuclear phagocyte subset in PBS-treated or DT-treated mice. Data is representative of two independent experiments. (i) The ratio of *Ccr2^−/−^*:wild-type (WT) cells within each of the indicated subset of C-LP mononuclear phagocyte from hematopoietic chimera reconstituted with a 1:1 ratio of WT and *Ccr2^−/−^* bone marrow. Data is representative of two independent experiments. (j) The geometric (geo) mean of acidified Zymosan pHrodo™ bioparticles within the indicated C-LP mononuclear phagocyte population after 1h incubation at 37°C. Fluorescence from cultures incubated at 4°C was subtracted. (k) The geometric mean of CFSE within the indicated C-LP mononuclear phagocyte population after 1h incubation with irradiated, CFSE labeled apoptotic thymocytes. CFSE fluorescence from cultures incubated at 4°C was subtracted. Data in (j,k) is combined from three independent experiments. Each dot represents pooled cells from 3-5 mice. (l) Percentage of IL-17^+^ OT-II CD4 T cells, and abundance of IL-17 protein (m) in co-cultures with CD26^+^ or CD26^−^ subsets within CD14^+^CD24^+^CD88^−^ APCs, analyzed as in Fig 1v,w. Data in (l,m) is combined from three independent experiments each containing sorted APCs from at least 5 mice. Each dot represents one experiment or replicates within an experiment. Error bars represent mean ± SEM. \**p* < 0.05, \*\**p* < 0.01, \*\*\**p* < 0.001, \*\*\*\**p* < 0.0001, n.s., not significant. (Statistical analysis for (i-k) was performed using one-way ANOVA with Tukey’s post hoc test. All other analyses were performed using unpaired Student’s t test).

Since CD26 expression resolved CD14^+^CD24^+^CD88^−^ APCs into two populations we sorted these into CD26^+^ and CD26^−^ subsets and analyzed their cellular appearance. The CD26^+^ subset had a dendritic appearance and in comparison to the known cDC subsets, most closely resembled CD103^−^CD11b^+^ DCs in size and slight vacuolar appearance (Fig 2d). By contrast, the CD26^−^ subset was highly vacuolar and did not have dendrites, consistent with the phenotype of intestinal macrophages, albeit the latter population was larger than the former.

Based on these cellular appearances we hypothesized the CD26^+^ subset belonged to the DC lineage, and the CD26^−^ subset belonged to the macrophage lineage. We undertook several tests to establish these lineage identities. First, we quantified their abundance in *Flt3l^−/−^* mice, since FLT3L is an essential hematopoietin for cDCs but is dispensable for macrophages^38–40^. Consistent with the DC-specific requirements for FLT3L, all cDC subsets were reduced 60-90% in *Flt3l^−/−^* mice (Fig 2e). Like cDCs, the CD26^+^ putative DC subset within CD14^+^CD24^+^CD88^−^ APCs also decreased by 50-80% in *Flt3l^−/−^* mice, suggesting that these required FLT3L and belonged to the DC lineage. By contrast, the CD26^−^ putative macrophage subset within CD14^+^CD24^+^CD88^−^ APCs was of equal abundance in *Flt3l^−/−^* and control mice, an outcome consistent with that of conventional macrophages and indicative of independence from FLT3L (Fig 2e,f).

Second, we tested for expression of two key lineage discriminating genes that are disparately expressed by macrophages and cDCs: 1) *Lyz2* (Lysozyme M), an antimicrobial enzyme most highly expressed by macrophages^41^, and 2) *Zbtb46*, a transcription factor expressed by all cDCs, but excluded in macrophages^42,43^. Consistent with their macrophage-like appearance, the CD26^−^CD14^+^CD24^+^CD88^−^ putative macrophage subset expressed robust *Lyz2*, at levels significantly higher than cDCs, and comparable to that of conventional macrophages (Fig S6, see Table 1 for all ANOVA comparisons). By contrast, the CD26^+^CD14^+^CD24^+^CD88^−^ putative DC subset expressed minimal *Lyz2*, and expression was not significantly different from cDCs. All cDCs expressed high levels of the cDC-specific transcription factor *Zbtb46*^42,43^, and although expression appeared reduced in both CD26^+^ and CD26^−^ subsets of CD14^+^CD24^+^CD88^−^ APCs, this did not reach significance. *Zbtb46* is a well-documented cDC specific gene and to clarify whether *Zbtb46* expression by CD14^+^CD24^+^CD88^−^ APCs was physiologically relevant we used *Zbtb46*-DTR hematopoietic chimera, wherein the administration of diphtheria toxin results in selective ablation of *Zbtb46*-expressing cells^42^. Indeed, diphtheria toxin reduced the numbers of cDCs and also CD26^+^CD14^+^CD24^+^CD88^−^ APCs by at least 65%, indicating *Zbtb46* expression by these populations was physiologically relevant (Fig 2g,h). By contrast, the CD26^−^CD14^+^CD24^+^CD88^−^ putative macrophage subset was only marginally reduced following diphtheria toxin (1.4 fold), and in this way these behaved similar to conventional macrophages in their resilience against *Zbtb46*-specific ablation. Thus, the putative CD26^+^CD14^+^CD24^+^CD88^−^ DC subset expresses *Zbtb46* at physiologically relevant levels, whereas the CD26^−^CD14^+^CD24^+^CD88^−^ putative macrophage subset did not.

Third, we tested the cell-intrinsic requirements for CCR2, a chemokine receptor that supports monocyte egress from bone marrow and is required to support population abundance of intestinal macrophages^13,34,44^. While macrophages are highly CCR2-dependent, CCR2 is also expressed and required for ~50% of CD103^−^CD11b^+^ cDCs, although the reasons underpinning this requirement remains unknown^15^. Both CD26^+^ and CD26^−^ populations within CD14^+^CD24^+^CD88^−^ APCs expressed uniformly high levels of CCR2 (Fig S5), suggesting that these novel APCs were CCR2-dependent. To test this, we generated mixed hematopoietic chimera using a mixture of bone marrow cells from WT (CD45.1) and *Ccr2^−/−^* (CD45.2) donors. To focus our analysis on mononuclear phagocytes originating from the adoptively transferred bone marrow we performed adoptive transfers into irradiated mice that co-expressed CD45.1 and CD45.2. Following immune reconstitution we identified host-derived cells as CD45.1^+^CD45.2^+^, excluded these from the analysis, and proceeded to evaluate each mononuclear phagocyte subset for their relative composition of *Ccr2^−/−^* and WT cells. As expected, intestinal macrophages derived from adoptively transferred bone marrow were highly CCR2-dependent, and essentially no macrophages developed from *Ccr2^-/-^* bone marrow (Fig 2i). By contrast, among cDCs only the CD103^−^CD11b^+^ subset had significant requirements for CCR2, a finding consistent with the fact that ~50% of these were CCR2^+^ and these are known to be CCR2-dependent^15^. The CD26^+^ and CD26^−^ subsets within CD14^+^CD24^+^CD88^−^ APCs were also highly CCR2-dependent, with <5% of these originating from *Ccr2^−/−^* bone marrow. Taken together with their appearance, physiologic expression of *Zbtb46* and disparate requirements for FLT3L, our results thus far suggest that the CD26^−^ subset within CD14^+^CD24^+^CD88^−^ APCs is a CCR2-dependent macrophage subset, and the CD26^+^ subset a CCR2-dependent DC subset.

Fourth, we tested phagocytosis, a function that is most robust in intestinal macrophages and is a validated measure to discriminate cellular functions of intestinal DCs and macrophages^20,45–47^. We chose zymosan for these tests since we found all CD14^+^CD24^+^CD88^−^ APCs expressed the zymosan-binding receptor Dectin-1 and also, that the CD26^+^ putative DC subset expressed the highest amount of Dectin-1 in comparison to all other intestinal mononuclear phagocytes (Fig S6). We used zymosan conjugated to a pH-sensitive fluorescent indicator (Zymosan pHrodo™) in which fluorescence is linked to acidification of zymosan-containing vesicles. Phagocytic uptake into acidified vesicles was robust in conventional macrophages, consistent with the functional specialization characteristic of intestinal macrophages (Fig 2j). Phagocytosis/acidification by the CD26^−^CD14^+^CD24^+^CD88^−^ putative macrophage subset was less than conventional macrophages but was nevertheless very robust, with most cells in this subset showing a positive signal (Fig S6). In comparison, phagocytic update of zymosan into acidified vesicles was low in both cDCs and the CD26^+^CD14^+^CD24^+^CD88^−^ putative DC subset, with no significant difference between the two. We also tested phagocytic uptake of apoptotic cells, a robust function of intestinal macrophages^20,45,46^. Phagocytic uptake of apoptotic cells was highest in conventional macrophages and CD26^−^CD14^+^CD24^+^CD88^−^ putative macrophages (Fig 2k, Fig S6). These two subsets had similar functional ability to uptake apoptotic cells and also, the functions of both subsets were significantly higher than that of cDCs and the CD26^+^CD14^+^CD24^+^CD88^−^ putative DC subset. Taken together, these data indicate that the CD26^−^CD14^+^CD24^+^CD88^−^ putative macrophage subset was highly phagocytic, a function consistent with their vacuolar appearance. By contrast, the CD26^+^CD14^+^CD24^+^CD88^−^ putative DC subset was minimally phagocytic, with functional ability comparable to that of cDCs.

Last, we tested the APC functions of CD26^+^ and CD26^−^ subsets within CD14^+^CD24^+^CD88^−^ populations and found Th17-inducing APC functions belonged to the CD26^+^ subset alone (Fig 2l,m). Taken together, the vacuolar or dendritic appearance, phagocytic potential, requirements for FLT3L or CCR2, *Zbtb46* expression, and abilities to activate naïve T cells classify CD14^+^CD24^+^CD88^−^ APCs into two novel subsets of CCR2-dependent mononuclear phagocytes: a CD26^+^CX3CR1^int/neg^ subset that belongs to the cDC lineage, and a CD26^−^ subset that belongs to the macrophage lineage. Remarkably, unlike other intestinal macrophages, CD26^−^CD14^+^CD24^+^CD88^−^ macrophages do not express CD88, CD64, or CX3CR1.

### CD26^−^CD14^+^CD24^+^CD88^−^ macrophages are an atypical subset with *Il10*^low^ and *Il6*^high^ gene expression

Having identified two new mononuclear phagocyte subsets with high specificity for the colon we next set out to address their gene expression. Among cytokine genes we focused on IL-10, IL-6, IL-23a (p19), and IL-12b (p40), since these potently influence APC functions and previous studies indicated that these were most disparately expressed by the different subsets of intestinal mononuclear phagocytes^6,10,48,49^. In particular, intestinal macrophages have a well-established *Il10*^high^ phenotype and this is thought to play a major role in intestinal homeostasis and macrophage-linked APC functions^16,49–52^. Although we predicted CD26^−^CD14^+^CD24^+^CD88^−^ macrophages would be *Il10*^high^, we were surprised to find that these expressed very little *Il10*, less than 8-fold that of conventional macrophages (Fig 3a; see Table 1 for all ANOVA comparisons). Another surprising distinction was for *Il6*—not only did CD26^−^CD14^+^CD24^+^CD88^−^ macrophages express the highest amount of *Il6* in comparison to all other subsets tested, but also, robust *Il6* expression was in stark contrast to that of conventional macrophages, which surprisingly expressed little to no IL-6 mRNA (Fig 3b). *Il6* was also expressed by DCs and although there was no statistical difference between the different DC subsets, expression trended higher in subsets with Th17-inducing APC functions: CD103^+^CD11b^+^ DCs, CD103^−^CD11b^+^ DCs, and also CD26^+^CD14^+^CD24^+^CD88^−^ DCs. *Il12b* and *Il23a* expression appeared DC specific since neither conventional macrophages or CD26^−^CD14^+^CD24^+^CD88^−^ macrophages expressed these cytokine genes (Fig 3c,d). All DC subsets expressed *Il12b*, whereas *Il23a* was most highly expressed by Th17-inducing DCs including CD26^+^CD14^+^CD24^+^CD88^−^ DCs, although these appeared to express lower (but not statistically significant) *Il12b* and *Il23a* as compared to the other Th17-inducing DC subsets.

**Figure 3.**
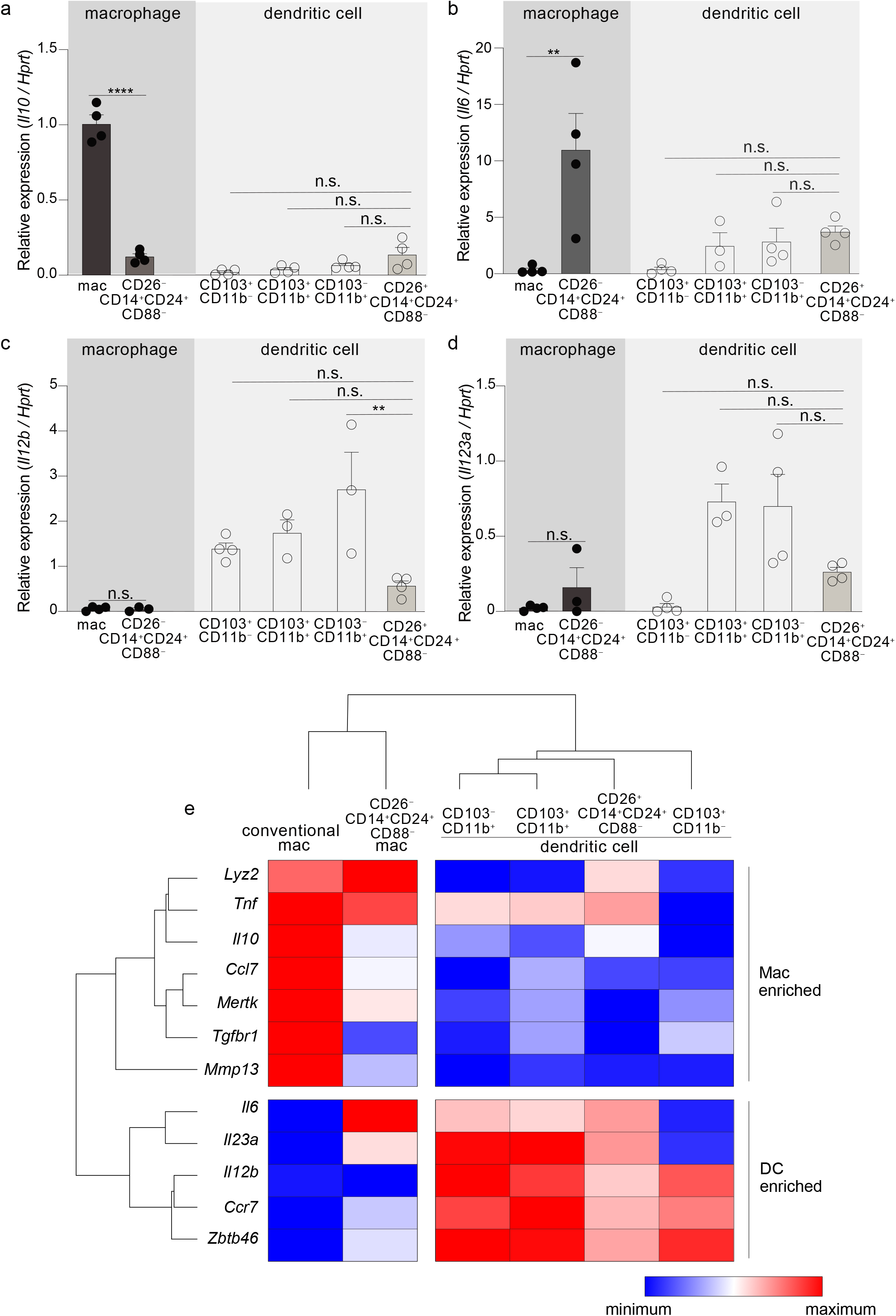
CD26^−^CD14^+^CD24^+^CD88^−^ macrophages are an atypical macrophage subset that expresses very little IL-10, and high levels of IL-6. Expression of *Il10* (a), *Il6* (b), *Il12b* (c), and *Il23a* (d) in the indicated subset of C-LP mononuclear phagocyte. Each data point represents expression in sorted cells pooled from at least 5 mice. (e) Unsupervised, hierarchical cluster analysis of Euclidian distance between the indicated C-LP mononuclear phagocyte subset based on the relative expression of the indicated gene. Data within each gene set is relative to *Hprt* and normalized to the average expression among all cell types tested. Error bars represent mean ± SEM. \*\**p* < 0.01, \*\*\*\**p* < 0.0001, n.s., not significant (one-way ANOVA with Tukey’s post hoc test).

To further expand upon these subset-specific patterns of gene expression we combined our analysis of cytokine gene expression with our analysis of lineage-differentiating genes (Fig S6) and performed unsupervised hierarchical clustering analysis to determine how gene expression aligned relationships between novel colon-specific APCs, cDCs and conventional macrophages. We also tested *Tnf* genes linked to the function and identity of DCs (*Ccr7, Zbtb46*), and genes linked to the function and identity of intestinal macrophages (*Lyz2, Mmp13, Tgfbr, Mertk, Ccl7*)^16,20,45^. Unsupervised hierarchical clustering analysis of these datasets distinguished CD26^+^CD14^+^CD24^+^CD88^−^ DCs and CD26^−^CD14^+^CD24^+^CD88^−^ macrophages as distally related populations, with the former population aligning with cDCs and the latter aligning with conventional macrophages (Fig 3e, see Table 1 for all ANOVA comparisons). Unsupervised hierarchical clustering analyses also revealed a relationship linkage between CD26^+^CD14^+^CD24^+^CD88^−^ DCs, CD103^−^CD11b^+^ and CD103^+^CD11b^+^ DCs, an outcome that may reflect the Th17-inducing APC functions shared by these subsets. There was also a close relationship between CD26^−^CD14^+^CD24^+^CD88^−^ macrophages and conventional macrophages, with both subsets being very distally related to DCs. CD26^−^CD14^+^CD24^+^CD88^−^ macrophages and conventional macrophages shared high expression of *Lyz2* and *Tnf*, but differed in the relative intensity of expression of other genes associated with conventional intestinal macrophages, including *Tgfbr1, Il6* and *Il10*. Taken together with our functional data and assessment of lineage-specific developmental requirements, unsupervised hierarchical clustering support lineage assignment of CD26^+^ and CD26^−^ colon-specific APCs as DCs and macrophages, respectively, and also highlight the unique gene expression profile of colon-specific CD26^−^CD14^+^CD24^+^CD88^−^ macrophages.

### Colon-specific DCs and macrophages require the transcription factor IRF4 and are absent from *Irf4*-cko mice

Given that CD26^+^CD14^+^CD24^+^CD88^−^ DCs possessed robust Th17-inducing APC functions we hypothesized that these would express IRF4. This transcription factor is universally expressed by all other known Th17-inducing DC subsets and is also required for their development and/or survival^14,15,25^. Consistent with our hypothesis CD26^+^CD14^+^CD24^+^CD88^−^ DCs expressed robust levels of IRF4 protein, with an MFI nearly equivalent to the two established IRF4^+^ DC subsets in the intestine: CD103^+^CD11b^+^ (<1.2 fold difference) and CD103^−^CD11b^+^ DCs (<1.4 fold difference) (Fig 5a and Fig S8)^14,15,25^. IRF4 protein was very weakly expressed by CD26^−^CD14^+^CD24^+^CD88^−^ macrophages, although expression appeared heterogeneous and the MFI of some cells within this macrophage population partially overlapped with that of Th17-inducing DCs (Fig 5a).

**Figure 5.**
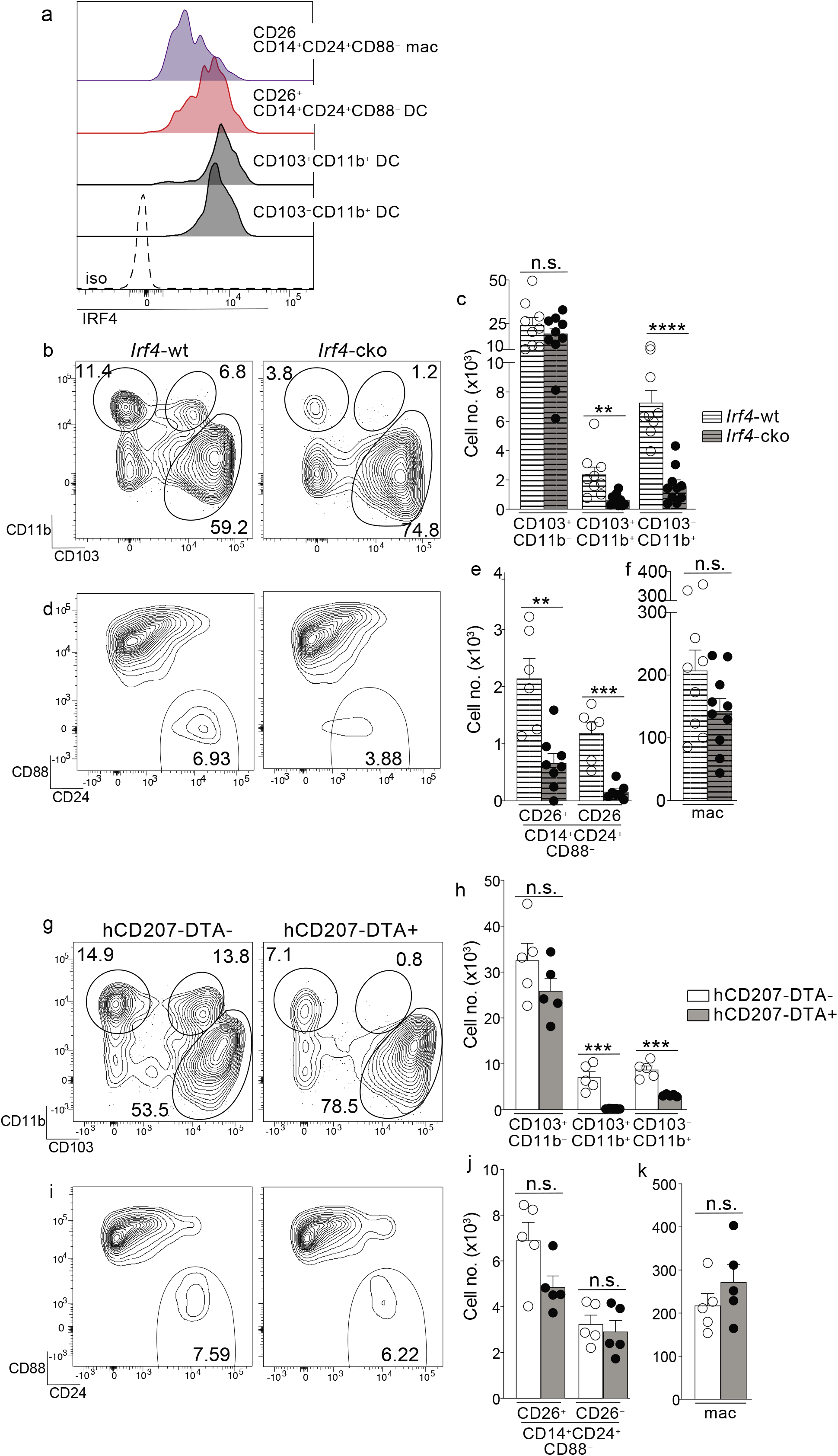
Colon-specific CD14^+^CD24^+^CD88^−^ DCs and macrophages are absent from *Irf4*-cko mice yet persist in hCD207-DTA mice. (a) Representative histogram of intracellular IRF4 protein in the indicated subset of C-LP mononuclear phagocyte. Iso, isotype control antibody staining. Data is representative of three separate experiments. (b-f) C-LP mononuclear phagocytes were analyzed from co-housed littermate *Irf4*-wt and *Irf4*-cko mice. Representative plot gated on total DCs as in Fig S1 (b) and the absolute cell number (c) of the indicated cDC subset. Representative plot gated as in Fig 1u of the abundance of CD14^+^CD24^+^CD88^−^ APCs (d) and the absolute cell number (e) of CD26^+^CD14^+^CD24^+^ CD88^−^ DCs and CD26^−^CD14^+^CD24^+^ CD88^−^ macrophages. (f) Absolute cell number of conventional macrophages from *Irf4*-wt or *Irf4*-cko mice. Data in (b-f) was combined from three independent experiments. (g-k) C-LP mononuclear phagocytes from co-housed littermate hCD207-DTA^+^ or hCD207-DTA^−^ mice were analyzed as in (b-f). Data are representative of three independent experiments. Error bars represent mean ± SEM. \*\**p* < 0.01, \*\*\**p* < 0.001, \*\*\*\**p* < 0.0001, n.s., not significant (unpaired Student’s t test).

Given these results, we tested whether IRF4 was required for the population abundance of colon-specific APCs. Since colon-specific DCs and macrophages were CD11c^+^ (Fig S3) we used *Irf4*-floxed x *Cd11c*-cre mice (*Irf4*-cko mice) to simultaneously test the requirements for IRF4 in multiple subsets of mononuclear phagocytes. Consistent with previous findings CD103^+^CD11b^+^ and CD103^−^CD11b^+^ cDCs were numerically reduced >70% in *Irf4*-cko mice, indicating that IRF4 was required for development and/or survival of these Th17-inducing DCs (Fig 5b,c)^14,25^. Importantly, CD14^+^CD24^+^CD88^−^ APCs were also markedly reduced in *Irf4*-cko mice, with numbers of CD26^+^ DCs within this population reduced >70% (Fig 5d,e). These data indicate that this novel colon-specific DC subset was highly IRF4-dependent.

While a requirement for IRF4 is well-established for Th17-inducing DCs, we were surprised to find that CD26^−^CD14^+^CD24^+^CD88^−^ macrophages were also numerically reduced >85% in *Irf4*-cko mice, suggesting a requirement for IRF4 in these atypical macrophages. (Fig 5e). Since the vast majority of CD26^−^CD14^+^CD24^+^CD88^−^ macrophages had little IRF4 protein (Fig 5a), this outcome may suggest that these low levels of IRF4 were nevertheless important for development and/or survival of this subset. Alternatively, it is possible that peak IRF4 expression occurred at an earlier stage of development that was not captured in our analysis. Although the mechanism underpinning the requirement for IRF4 remains to be determined, it was clear that depletion of CD26^−^CD14^+^CD24^+^CD88^−^ macrophages in *Irf4*-cko mice was in stark contrast to conventional macrophages, since these were numerically similar in both *Irf4*-cko and control mice (Fig 5f). Taken together our data indicate that colon-specific CD14^+^CD24^+^CD88^−^ DCs and macrophages require the transcription factor IRF4.

### Colon-specific DCs and macrophages persist in hCD207-DTA mice and are independent of other IRF4-dependent APCs

Given our results showing that *Irf4*-cko mice lacked colon-specific CD14^+^CD24^+^CD88^−^ DCs and macrophages, as well as other IRF4-dependent APCs (CD103^+^CD11b^+^ and CD103^−^CD11b^+^ DCs), we sought to determine whether the novel populations we described here were derived from other IRF4-dependent APCs, or if they were autonomous. To test this we sought a strategy wherein numbers of colon-specific CD14^+^CD24^+^CD88^−^ APCs remained intact, while all other IRF4-dependent APCs were absent. To this end we tested hCD207-DTA mice, where the human CD207 promoter drives intracellular expression of diphtheria toxin and consequentially, subset-specific ablation of mononuclear phagocytes that transcribe the hCD207-DTA transgene^26,53^. In this fashion, CD103^+^CD11b^+^ and CD103^−^CD11b^+^ DCs, which are absent from *Irf4*-cko mice, are also ablated in hCD207-DTA mice^10,26^. If colon-specific CD14^+^CD24^+^CD88^−^ APCs were autonomous populations from CD103^+^CD11b^+^ and CD103^−^CD11b^+^ DCs, we reasoned that these would not transcribe the hCD207 transgene and thus, would not be ablated in hCD207-DTA mice. We examined hCD207-DTA mice and co-housed littermate control mice to test this directly.

As expected, hCD207-DTA mice were markedly deficient in colonic CD103^+^CD11b^+^ and CD103^−^CD11b^+^ DCs, with numerical abundance of each population decreased 96% and 65%, respectively (Fig 5g,h). In stark contrast to these, colon-specific CD14^+^CD24^+^CD88^−^ DCs and macrophages were intact in hCD207-DTA mice, with no significant difference found between hCD207-DTA mice and co-housed littermate controls (Fig 5i,j). Numbers of conventional macrophages were also similar between the two mouse strains (Fig 5k). The persistence of colon-specific CD14^+^CD24^+^CD88^−^ DCs and macrophages in hCD207-DTA mice indicates that these APC subsets did not transcribe the hCD207 transgene, and thus were distinct populations from CD103^+^CD11b^+^ and CD103^−^CD11b^+^ DCs, which were ablated from these same mice. Moreover, that CD103^+^CD11b^+^ and CD103^−^CD11b^+^ DCs were absent from hCD207-DTA mice strongly suggests that colon-specific CD14^+^CD24^+^CD88^−^ APCs (which persist in hCD207-DTA mice) arise independent from other IRF4-dependent mononuclear phagocytes and are not “alternative states” of any other IRF4-dependent APC subset. Thus, hCD207-DTA mice retain colon-specific CD14^+^CD24^+^CD88^−^ DCs and macrophages, but other IRF4-dependent APCs are absent.

### CD14^+^CD24^+^CD88^−^ APCs are required for Th17 immunity in the colon

Our results above indicated that colons of *Irf4*-cko and hCD207-DTA mice contained distinct APCs: *Irf4*-cko mice lacked all Th17-inducing APCs, including colon-specific APCs (Fig 5b-f), whereas hCD207-DTA mice retained colon-specific APCs, yet lacked other Th17-inducing APCs (Fig 5g-k). We leveraged these findings and evaluated Th17 cells in *Irf4*-cko and hCD207-DTA mice to determine if colon-specific APCs were required for Th17 immunity *in vivo*.

We first evaluated *Irf4*-cko mice, which lack all Th17-inducing APCs, including colon-specific APCs. Consistent with a requirement for IRF4-dependent APCs to support intestinal Th17 immunity^14,25^, colon-resident Th17 cells were markedly reduced in *Irf4*-cko mice, with numbers >4.5 fold fewer than that of co-housed littermate controls (Fig 6a-c). Th17 cells were also significantly reduced in small intestine of *Irf4*-cko mice, confirming that IRF4^+^ APCs were essential for Th17 immunity in both small intestine and colon (Fig 6d,e).

**Figure 6.**
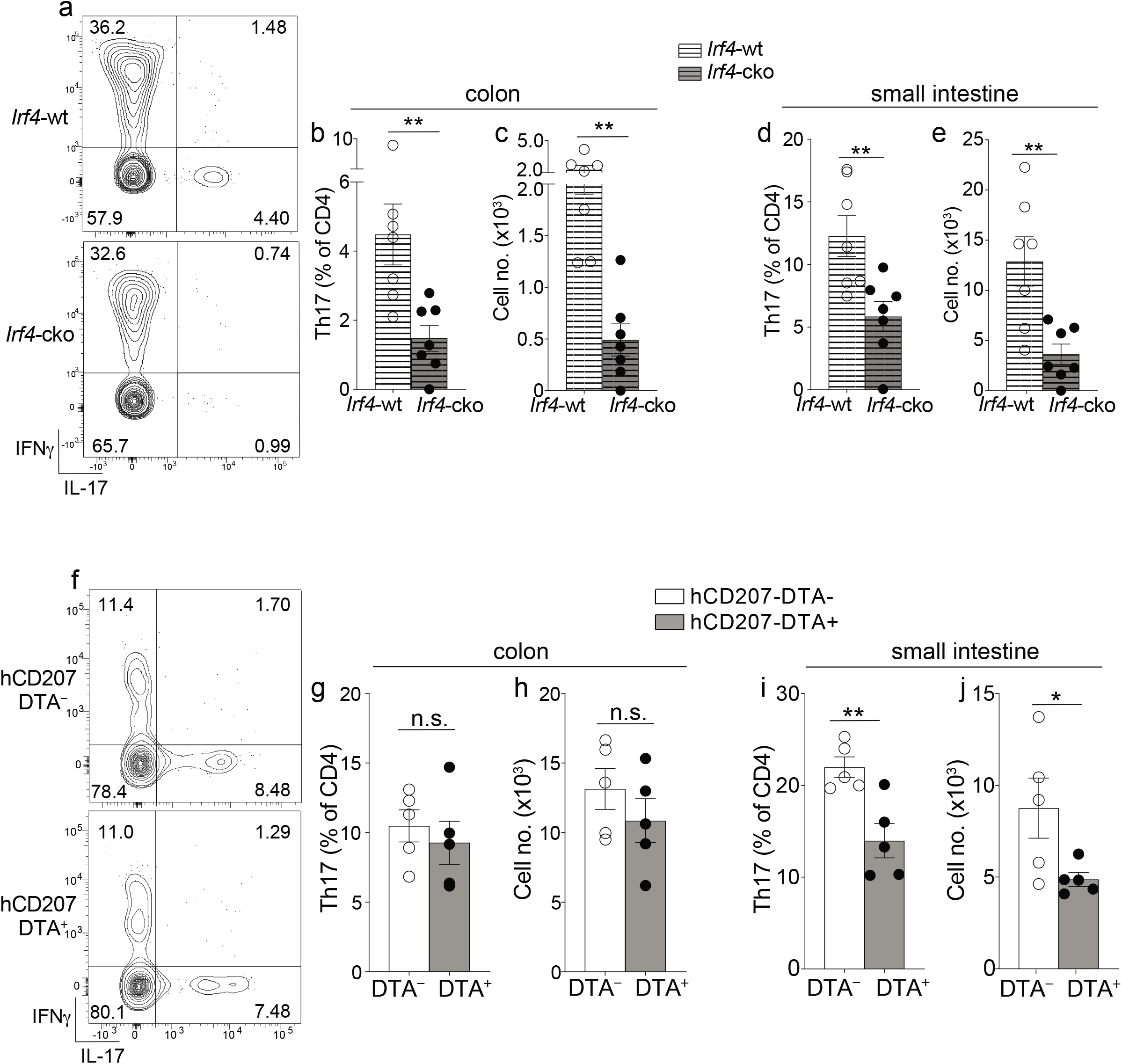
Colon-specific CD14^+^CD24^+^CD88^−^ APCs are required for Th17 immunity in the colon. (a-e) C-LP or SI-LP CD4 T cells were analyzed from co-housed littermate *Irf4*-wt and *Irf4*-cko mice. (a) Representative plot of C-LP CD4 T cells from mice of the indicated genotype analyzed for IFNγ and IL-17 production. The percent (b) and absolute cell number (c) of C-LP Th17 cells from all mice tested. The percent (d) and absolute cell number (e) of SI-LP Th17 cells in *Irf4*-wt and *Irf4*-cko mice analyzed in (b,c). Data in (b-e) was combined from three independent experiments. (f-j) C-LP and SI-LP CD4 T cell populations were analyzed from co-housed littermate hCD207-DTA^−^ and hCD207-DTA^+^ mice as in (a-e). Representative plot of C-LP CD4 T cells (f), the percent (g) and absolute cell number (h) of C-LP Th17 cells, and the percent (i) and absolute cell number (j) of SI-LP Th17 cells. Data are representative of three independent experiments. Error bars represent mean ± SEM. \**p* < 0.05, \*\**p* < 0.01, n.s., not significant (unpaired Student’s t test).

In stark contrast to *Irf4*-cko mice, colon Th17 cells were of normal abundance in hCD207-DTA mice, and both the percent and cell number of colon-resident Th17 cells were similar between hCD207-DTA mice and co-housed littermate controls (Fig 6f-h). Despite the persistence of Th17 cells in colon, Th17 cells in small intestine of hCD207-DTA mice were markedly reduced (Fig 6i,j). Depletion of Th17 cells from small intestine of hCD207-DTA mice is consistent with the loss of CD103^+^CD11b^+^ and CD103^−^CD11b^+^ DCs from this organ (Fig S9 and^26^). That Th17 cells were lost from small intestine but retained in the colon of hCD207-DTA mice clearly indicates that these organs have distinct APC requirements for Th17 immunity. Based on the discoveries we report here, evidence suggests the basis for this organ-specific distinction is due to the fact that CD103^+^CD11b^+^ and CD103^−^CD11b^+^ DCs are the only Th17-inducing APCs in small intestine, whereas colon also contains a novel population of CD14^+^CD24^+^CD88^−^ APCs. Since colon-specific APCs persist in hCD207-DTA mice, yet are absent in *Irf4*-cko mice, our results further suggest that CD14^+^CD24^+^CD88^−^ APCs are both required and sufficient for Th17 immunity in the colon (See Fig S10 for graphical abstract). Colon-specific residence of CD14^+^CD24^+^CD88^−^ APCs likely underscores their dedicated role in Th17 immunity exclusively in this intestinal organ.

## Discussion

The magnitude and diversity of antigens encountered by the intestinal immune system is beyond compare. Understanding how this demand is met requires precise identification of the cellular constituents of intestine-resident immune populations. To this end, our initial goal was to disambiguate MHC-II^+^CD11c^+^/CD11b^+^ APCs that co-expressed CD14 and CD24. Although these were few in small intestine, colon contained many CD14^+^CD24^+^ APCs and we surprisingly revealed that this population contained two novel mononuclear phagocyte subsets: a Th17-inducing cDC subset and an unconventional, CX3CR1^−^*Il10*^low^ macrophage subset. Neither of these populations were found in small intestine. Each intestinal organ has unique immunological demands and our studies reveal that organ-specific mononuclear phagocyte populations arise to meet these demands. Thus, specific cell subsets within the mononuclear phagocyte network have dedicated organ-specific roles in intestinal immunity.

The implementation of CD88 for subsetting intestine-resident immune populations proved to be a remarkable asset for our discovery of colon-specific mononuclear phagocytes, and this utility for CD88 is a key advance from this study. Among MHC-II^+^ APCs CD88 was previously reported to be expressed on macrophages and monocyte-derived DCs^36^. That same report demonstrated that CD88 could distinguish MHC-II^+^ macrophages and monocyte-derived DCs populations from cDCs (which are CD88^negative^), and that this utility for CD88 was applicable to many tissues and across mouse strains^36^. Our findings reinforce CD88’s utility for subsetting mononuclear phagocytes in the intestine and we find CD88’s best asset to be its high expression by all intestinal macrophages, with notable exception of the atypical colon-specific macrophage we identify here. Importantly, we also show that CD88 is robustly upregulated in transitioning monocytes undergoing development into macrophages during the “monocyte-waterfall” and that by virtue of this high level of expression, CD88 enables accurate identification of still-developing cells committed to the macrophage lineage. Currently, identification of transitioning monocytes requires that these express Ly6C, a marker that is progressively downregulated during the monocyte-waterfall^13,35^. That these concomitantly upregulate CD88 enables this marker to be used to identify still-developing transitioning monocytes that have downregulated Ly6C but have yet to acquire the mature macrophage phenotype. In this way the inclusion of CD88 to study the “monocyte-waterfall” should provide new insight into the incremental steps involved in the transition of monocyte to macrophage in the intestine, a process which is poorly defined.

We also demonstrate the importance of using CD88 together with CD14 to properly subset intestinal mononuclear phagocytes. Previous work identified intestinal macrophages using either CD14 alone or CD64 alone and these markers were not only thought to be co-expressed by intestinal macrophages, but it was also thought that these could be used interchangeably to distinguish intestinal macrophages and DCs^11,13–15,20^. Our findings now make clear that CD14 and CD64 are not interchangeable, nor always co-expressed by macrophages, nor are they excluded in all intestinal cDC subsets, since we identified a macrophage subset and a cDC subset that both share a CD14^+^CD64^negative^ phenotype. Both of these novel CD14^+^CD64^negative^ subsets are CD103^negative^ and thus, previous studies using CD64 alone to distinguish macrophages from DCs would result in an erroneous mixture of different cell subsets within the population designated as the CD103^−^CD11b^+^ DC subset—this mixture would contain CD103^−^CD11b^+^ DCs as well as the CD14^+^CD24^+^CD88^−^ DCs and macrophages we describe here. The erroneous mixture arising from the use of CD64 alone is corrected with the use of CD14 and indeed, if not for their differences for CD14 expression, CD103^−^CD11b^+^ DCs and colon-specific CD14^+^CD24^+^CD88^−^ DCs appear very similar. Both are CX3CR1^int/neg^, CCR2^+^, and both possess Th17-inducing APC functions. Since CD14^+^CD24^+^CD88^−^ DCs are the only CD14^+^ intestinal DC subset, it is possible that CD14 serves a unique function in these DCs, which are colon-specific. CD14 has well-described roles as the LPS co-receptor in innate immune signaling, and in this manner CD14 could influence how CD14^+^CD24^+^CD88^−^ DCs respond to microbiota and microbial pathogens. Unfortunately, assigning a functional role for CD14 in colon-specific DCs is not possible at this time since CD14 is required as a cell surface marker to distinguish this population, and this requirement creates a barrier to functional studies using cells from CD14-deficient mice. Exploration of a functional role for CD14 on colon-specific DCs awaits further studies that implement other unique identifying markers, once these are identified.

Colon-specific CD14^+^CD24^+^CD88^−^ DCs are a distinct DC subset, separate from the other IRF4-dependent DCs that reside in the intestine. Thus, while CD103^−^CD11b^+^ and CD103^+^CD11b^+^ DCs transcribe and are ablated by the hCD207-DTA transgene, CD14^+^CD24^+^CD88^−^ DCs do not. The persistence of this DC subset in hCD207-DTA mice is good evidence that CD14^+^CD24^+^CD88^−^ DCs are a *bona fide* DC subset and not an “alternate state” of any other. While our data make clear that CD14^+^CD24^+^CD88^−^ DCs do not originate from CD103^−^CD11b^+^ DCs or CD103^+^CD11b^+^ DCs, we cannot rule out the possibility that the latter two populations originate from CD14^+^CD24^+^CD88^−^ DCs. However, this relationship is unlikely given that CD103^−^CD11b^+^ DCs and CD103^+^CD11b^+^ DCs are present in both small intestine and colon, whereas CD14^+^CD24^+^CD88^−^ DCs are colon-specific. Transcriptional profiling of colon-specific CD14^+^CD24^+^CD88^−^ DCs should shed light on these questions and is a high priority for future work given our results showing the dedicated and essential role for these in Th17 immunity in the colon.

That colon and small intestine have distinct APC requirements is one of the most notable findings of our report, since it was previously assumed that CD103^−^CD11b^+^ and CD103^+^CD11b^+^ DCs were essential for Th17 immunity in both intestinal organs. This assumption was largely based on reports showing that mice lacking CD103^−^CD11b^+^ and CD103^+^CD11b^+^ DCs have reduced numbers of Th17 cells in the small intestine^10,25,26^—pointedly, colon was not investigated in those previous reports. Here we show that in stark contrast to small intestine, the absence of CD103^−^CD11b^+^ and CD103^+^CD11b^+^ DCs from the colon of hCD207-DTA mice has no negative impact to colon-resident Th17 cells. Th17 immunity in colon instead requires CD14^+^CD24^+^CD88^−^ APCs, which are colon-specific.

Understanding why CD14^+^CD24^+^CD88^−^ DCs express and require CCR2 is another priority, since CCR2 is best known as a chemokine receptor for monocyte-derived populations. Although CCR2-dependent, it is highly unlikely that monocytes give rise to CD14^+^CD24^+^CD88^−^ DCs, since these DCs express *Zbtb46* and are FLT3L-dependent. CD14^+^CD24^+^CD88^−^ DCs appear to be similar to CD103^−^CD11b^+^ DCs, which also express and require CCR2, yet belong to the conventional DC lineage^15^. Another unusual phenotype shared by CD103^−^CD11b^+^ and CD14^+^CD24^+^CD88^−^ DCs is that both express low levels of the monocyte/macrophage chemokine receptor CX3CR1. As a bulk population, CX3CR1^+^ cells support immunity to intestinal fungi, bacterial pathogens, and also support oral tolerance^54,55^. Dissecting out the individual roles of each CX3CR1^+^ subset, a population that includes macrophages, CD103^−^CD11b^+^ DCs and CD14^+^CD24^+^CD88^−^ DCs, is an important priority. Reasons underpinning why CX3CR1 and CCR2 are expressed by two cDC subsets in the intestine are unknown, although it should be noted that CCR2 and CX3CR1 are expressed by pre-DCs and these chemokine receptors support pre-DC homing to mucosal tissue^56,57^. Thus, a potential explanation as to why CCR2 and CX3CR1 are retained on CD103^−^CD11b^+^ and CD14^+^CD24^+^CD88^−^ DCs may be that these are derived from pre-DCs that fail to downregulate these markers during development in the colon. There may be a pre-DC subset committed to this developmental path since pre-DCs can be subsetted based on their commitment to becoming either an IRF8^+^ or IRF4^+^ DC^57^. Given that we show colon contains three different subsets of IRF4^+^ Th17-inducing DCs, it is possible that multiple pre-DC subsets are needed to meet the unique immunological demands of this intestinal organ.

One of the most surprising findings of our report is the identification of colon-specific macrophages whose phenotype and function deviate substantially from all other intestinal macrophages. Phenotypically, colon-specific CD14^+^CD24^+^ macrophages fail to express CD88, CD64 and also fail to express CX3CR1. Also, colon-specific macrophages are dependent on the transcription factor IRF4, a requirement that is unique and specific to this intestinal macrophage subset. Naturally occurring IRF4-dependent macrophages appear to be rare *in vivo* and other than the macrophage subset we report here, an *in vivo* requirement for IRF4 in macrophages has only been described for F4/80^low^MHC-II^+^CD226^+^ macrophages resident in the peritoneal and pleural cavity^58^. Although these macrophages are not CX3CR1^negative^, CX3CR1 expression is nevertheless downregulated during their development, and thus the IRF4-dependent process that gives rise to these CX3CR1^low^ macrophages from monocyte precursors may be similar to the developmental process that gives rise to the CX3CR1^negative^ macrophages in colon that we describe here. Since the majority of colon-specific macrophages do not express IRF4 protein, this would suggest that continuous expression of IRF4 is not required for the maintenance, survival or steady-state gene expression of CD14^+^CD24^+^CD88^−^ macrophages. Thus, the requirement for IRF4 must occur during development, presumably during a stage of transient IRF4 expression in transitioning monocytes. Monocytes are now also appreciated to exist in many subsets and it is possible that there is a dedicated subset that upregulates IRF4 to give rise to CD14^+^CD24^+^CD88^−^ macrophages^59^. While the developmental process of colon-specific macrophages needs to be further explored, our studies reveal the existence of two developmental paths for intestinal macrophages—an IRF4-independent path that gives rise to conventional CD88^+^CD64^+^CX3CR1^+^ macrophages, and an IRF4-dependent path that generates unconventional CD14^+^CD24^+^ macrophages that do not express CD88, CD64, or CX3CR1.

Surprisingly, this IRF4-dependent path gives rise to macrophages which are *Il10*^low^ and *Il6*^high^. In fact, IL-6 mRNA expression by colon-specific macrophages exceeded that of all other mononuclear phagocytes tested. IL-6 is essential for the instruction of Th17 cells and by providing robust IL-6 stimulation, combined with the lack of antagonistic IL-10, CD14^+^CD24^+^CD88^−^ macrophages could potently influence Th17 immunity. Although our data indicate that CD14^+^CD24^+^CD88^−^ macrophages cannot instruct Th17 development *in vitro*, it is possible that they do so *in vivo*, since intestinal macrophages are thought to be sufficient for Th17 immunity to the intestinal microbe segmented filamentous bacteria^60^. Even if CD14^+^CD24^+^CD88^−^ macrophages do not directly instruct naïve T cells, robust IL-6 expression by these likely collaborates with DC-mediated instruction of Th17 cells *in vivo*. This collaboration could occur in the colon itself since evidence suggests that lymph nodes are not required for the genesis of microbiota-reactive Th17 cells and that these can develop *in situ*, in intestinal lamina propria^23,24^. In this manner, CD14^+^CD24^+^CD88^−^ macrophages within the intestinal microenvironment could have major impact on the instruction of Th17 cells.

Finally, our findings reveal the existence of organ-specific factors that give rise to CD14^+^CD24^+^CD88^−^ macrophage and DC subsets that are uniquely colon specific. Identification of these factors and how they initiate the IRF4-dependent pathway for macrophage development, or the pathway that gives rise to CD14^+^ cDCs, is a top priority for future studies. Presumably these factors are absent or antagonized in the small intestine. Why colon requires additional subsets of mononuclear phagocytes in comparison to the small intestine is likely linked to the unique anatomy and function of these organs, as well as unique vulnerabilities to pathogens. Having identified novel colon-specific mononuclear phagocytes and demonstrating their essential function for Th17 immunity in the colon during steady-state, our findings here compel future studies to elucidate the responses of colon-specific mononuclear phagocytes in the setting of pathogen infection.

## Materials and methods

### Mice

*Irf4*-floxed, *Cd11c*-cre, *Zbtb46*-DTR, *Cx3cr1*-GFP, *Ccr2^−/−^* and hCD207-DTA transgenic mice have been described previously^26,42,61–63^. Those mice were purchased from The Jackson Laboratory and were housed under specific pathogen free conditions at Duke University. Animal protocols were approved by the Duke University Institutional Animal Care and Use Committee. *Flt3l^−/−^* mice were on a 129 background and were housed under SPF conditions at the Durham VA Medical Center. All mice were analyzed between 10-14 weeks of age.

### Intestinal lamina propria preparation

Colon, small intestine, and cecum lamina propria cells were prepared as previously described^10^. Briefly, intestinal tissue was cut longitudinally and luminal contents were removed by vigorous washing. Tissues were then cut into 0.2-0.3cm pieces and washed twice using HBSS/10mM HEPES/5mM EDTA/1.25% BSA/1mM DTT for 10 minutes, followed by another 10 min wash in HBSS/10mM HEPES/1.25%BSA. Tissues were digested in an enzyme mixture of Liberase™ (57.6 μg/ml) and DNase I (8 U/ml) in C-tubes (Miltenyi) for 30 min at 37°C with constant shaking, followed by dissociation using the GentleMACS™ Dissociator (Miltenyi). After digestion, cellular suspension was filtered and subjected to RBC lysis prior to analysis.

### Flow cytometry immune cell phenotyping and colonic lamina cell sorting

Freshly isolated lamina propria cells were stained with FcR block (Biolegend) and Live/dead^®^ fixable dead cell staining (Thermo Fisher) and gated with the following markers: Monocytes, (CD11b^+^CD24^−^Ly6C^+^IA/E^−^); transitioning monocytes, (CD11b^+^CD24^−^Ly6C^+^IA/E^+^); conventional macrophages, (CD11b^+^IA/E^+^CD14^+^CD24^−^Ly6C^−^); conventional DCs, (IA/E^+^CD11c^+^CD24^+^CD14^−^CD88^−^Ly6C^−^, followed by CD103 and CD11b to distinguish conventional DC subsets); novel colon-specific CD26^+^CD14^+^CD24^+^CD88 DCs, (IA/E^+^CD11b^+^CD11c^+^CD24^+^CD14^+^Ly6C^−^CD88^−^CD26^+^); novel colon-specific CD26^−^CD14^+^CD24^+^CD88^−^ macrophages, (IA/E^+^CD11b^+^CD11c^+^CD24^+^CD14^+^Ly6C^−^CD88^−^CD26^−^); CD4 T cells (CD45^+^TCRβ^+^CD4^+^). For intracellular cytokine staining, lamina propria cells were stimulated for 5 h with PMA (50 ng/ml) and ionomycin (500 ng/ml) in the presence of BD Golgi plug (BD Biosciences) followed by fixation and permeabilized using the BD Cytofix/Cytoperm kit. Flow cytometry was performed on a BD Fortessa X20 or BD Canto (BD Biosciences), and data analyzed using FlowJo software. To sort specific subsets of C-LP mononuclear phagocytes cells were gated as above and sorted by MoFlo Astrios Sorter (Beckman Coulter).

### Hematopoietic chimera

To generate *Zbtb46*-DTR bone marrow chimera, 10-week old CD45.1 mice were irradiated with two doses of 600cGy (X-RAD 320) with 3 h rest between doses. Irradiated mice were injected intravenously with 2×10^6^ bone marrow cells from *Zbtb46*-DTR mice (CD45.2). To generate WT and *Ccr2^−/−^* mixed hematopoietic chimera, 10-week-old CD45.1^+^CD45.2^+^WT mice were irradiated as above and reconstituted with a 1:1 mixture containing 2×10^6^ each of WT bone marrow (CD45.1) and *Ccr2^−/−^* (CD45.2) bone marrow. All hematopoietic chimera were subsequently treated or analyzed 8 weeks post reconstitution.

### Phagocytosis assays

Freshly isolated C-LP cells pooled from 3-5 mice were co-cultured in duplicate with pHrodo™ Zymosan Bioparticles™ (Life Technology) at 37°C or 4°C in complete RPMI medium for 1 h according to manufacturer’s instructions. Co-cultures at 4°C were analyzed in tandem and used as negative controls. Cell surface markers and pHrodo signal fluorescence was analyzed by flow cytometry and data reported as the geometric mean (37°C - 4°C) of the pHrodo signal. For phagocytosis of apoptotic thymocytes, mouse thymocytes were exposed to 254 nm UV-irradiation for 6 minutes (Stratagene Stratalinker) followed by labelling with 2 μM CFSE (Life Technologies) and subsequent culture for 24 h a 37°C in serum-free RPMI.

Thymocyte cultures were routinely 100% CFSE^+^ and ~50% Annexin V^+^. Apoptotic thymocytes were then co-cultured 10:1 with freshly prepared C-LP cells and incubated for 1 h at 37°C in 24-well ultra-low attachment plates (Corning). Separate cultures incubated at 4°C were used as controls. Cell surface markers and CFSE fluorescence in mononuclear phagocyte populations were analyzed by flow cytometry and data reported as the geometric mean (37°C - 4°C) of CFSE.

### Diphtheria toxin ablation of *Zbtb46*-expressing cells

*Zbtb46*-DTR hematopoietic chimera were injected intraperitoneally with 40 ng/g of diphtheria toxin (Sigma-Aldrich) or PBS as a negative control. C-LP cells were analyzed 24 h post injection.

### Microscopy

Sorted C-LP mononuclear phagocytes from 5-7 mice were centrifuged onto slides using the Cytospin 2 (Shandon). Slides were dried in a desiccation chamber overnight and stained with Fisher HealthCare™ PROTOCOL™ Hema 3™ Manual Staining System (Thermo scientific). Images were captured using a light microscopy (Zeiss Axio Imager Widefield fluorescence microscope) at 40x magnification and were processed by Image J.

### DC and OT-II T cell co-culture

Co-cultures were performed using sorted naïve OT-II T cells (CD4^+^TCRβV5.1/5.2^+^CD44^−^CD62L^+^CD25^−^) and sorted C-LP APCs, as described^10^. Briefly, these were co-cultured 1:1 in complete RPMI containing OVA peptide (5 mg/mL, InvivoGen) and recombinant TGFβ (1 ng/mL, PeproTech) for 96 h. To assay cytokine secretion cells were re-stimulated for 4 hours with plate bound anti-CD3 (5 mg/mL) and anti-CD28 (5 mg/mL) in the presence of BD Golgi plug. Cytokine production by re-stimulated OT-II CD4 T cells was analyzed by gating on live CD4^+^TCRβ^+^, followed by fixation and permeabilization using the BD Cytofix/Cytoperm kit followed by analysis of intracellular IFNγ and IL-17 protein. Total numbers of CD4 T cells were calculated from unstimulated cultures by flow cytometry using CountBright™ absolute counting beads (Thermo Fisher). IL-17 and IFNγ in supernatants were assayed by ELISA (Biolegend or R&D systems). For data output cell number and ELISA were normalized per 1,000 of each APC subset used for co-culture.

### Quantitative PCR

C-LP cells pooled from 5-7 mice were sorted into TriZol^®^ (Thermo Fisher) and cDNA was generated by using QuantiTect Reverse Transcription Kit (Qiagen). Quantitative PCR reactions were performed using TaqMan™ Gene Expression Master Mix (AppliedBiosystems) with the following TaqMan probe assays: *Il6* (Mm00446190_m1), *Il23a* (Mm00518984_m1), *Il12b* (Mm00434174_m1), *Il10* (Mm00439614_m1), *Zbtb46* (Mm00511327_m1), *Tgfbr1* (Mm00436964_m1), *Mertk* (Mm00434920_m1), *Ccl7* (Mm00443113_m1), *Lyz2* (Mm01612741_m1), *Tnf* (Mm00443258_m1), *Mmp13* (Mm00439491_m1), *Ccr7* (Mm01301785_m1), and *Hprt* (Mm03024075_m1). Hierarchical clustering of relative gene expression of quantitative PCR gene set was performed using web-based open source software Morpheus (cite: Morpheus, http://software.broadinstitue.org/morpheus). Briefly, relative gene expression was normalized to average expression among all indicated C-LP mononuclear phagocytes and then log2 transformed. Hierarchical clustering was performed using Euclidian distance metric and complete linkage method.

### Statistical analyses

Statistical analysis was performed using GraphPad Prism (GraphPad, San Diego, CA) to perform 2-tailed Student’s t test or one-way analysis of variance (ANOVA) with Tukey’s or Dunnett’s post hoc multiple comparisons test.

## Acknowledgments

This work was supported by NIH R01-AI45930 (awarded to G.E.H.). Gianna Elena Hammer is a Pew Biomedical Scholar and this work was additionally funded by the Pew Charitable Trusts.

## Author contributions

GEH conceived the study, acquired funding, and provided supervision. HH, MLJ, NY, MH, BEF, and ER executed experiments and performed data analysis. ERH supervised statistical analysis. NPR, JRP, JJZ, DW, GAT, MDF, DEK, DNC, VC and SDC provided reagents. GEH wrote the manuscript, which was edited by HH and MLJ.

## Supplemental figure legends

**Figure S1.**
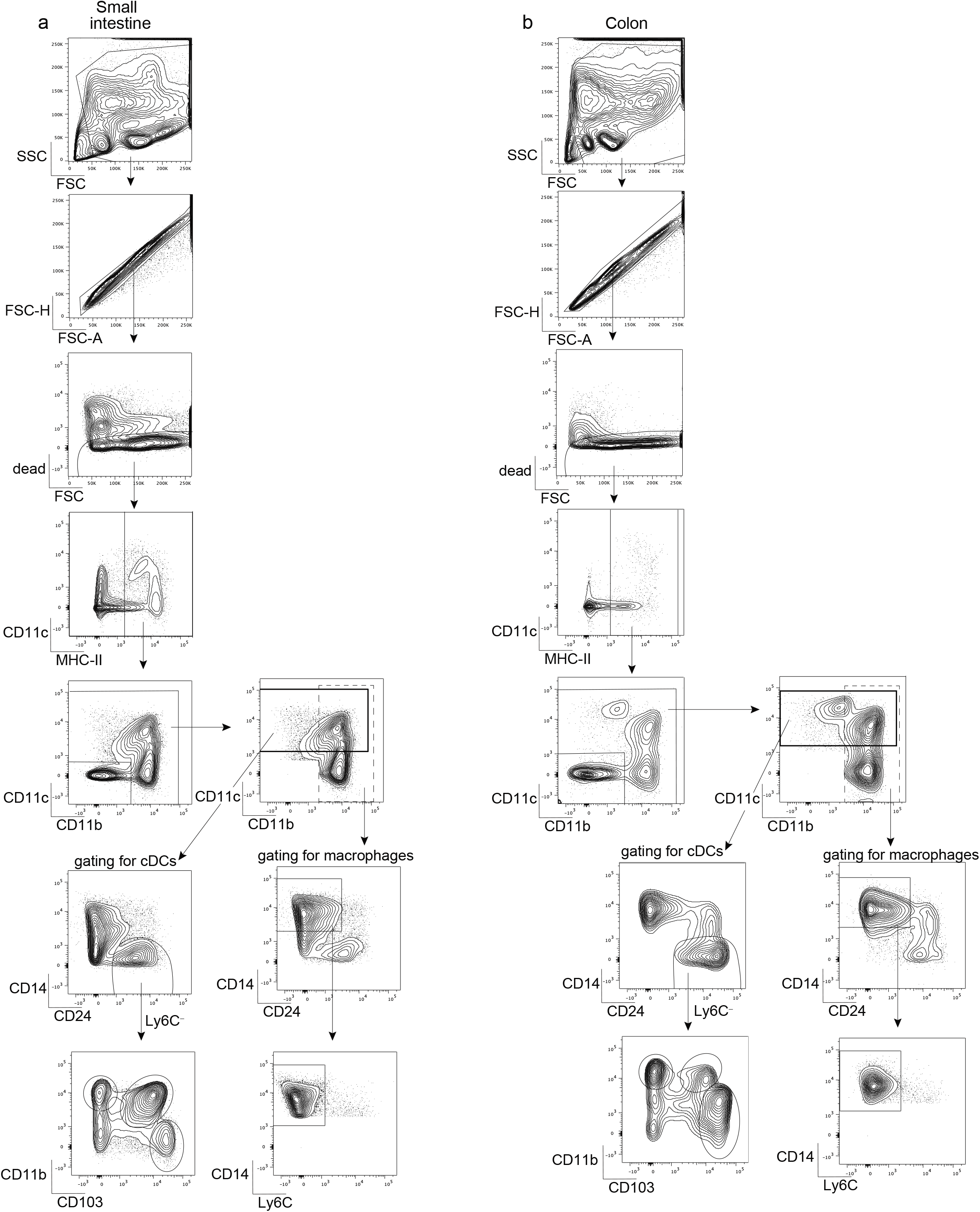
**Related to Figure 1** (a-b) Gating strategy to identify CD103^−^CD11b^+^, CD103^+^CD11b^−^, and CD103^+^CD11b^+^ cDC subsets and conventional macrophages in lamina propria of small intestine (a) and colon (b).

**Figure S2.**
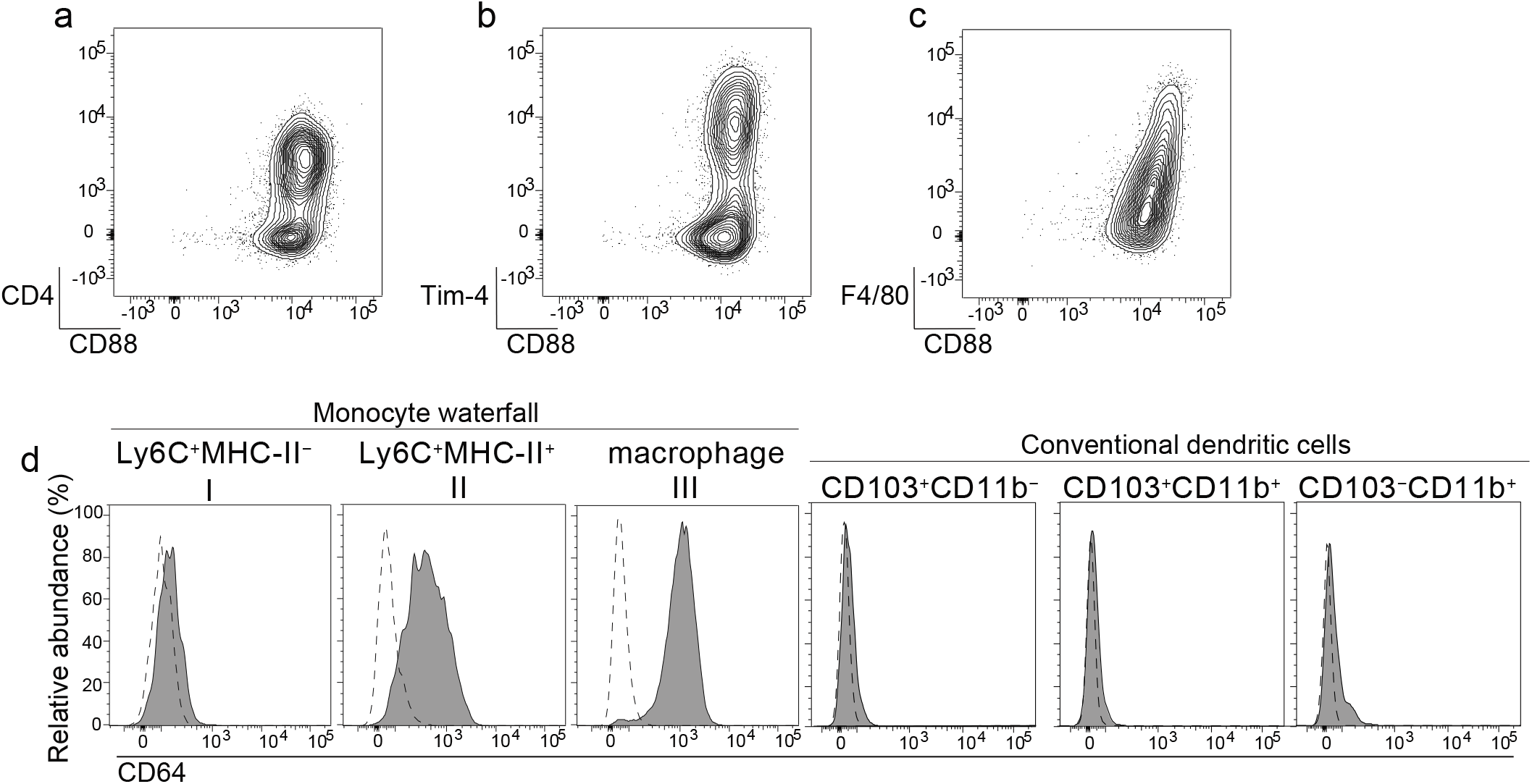
**Related to Figure 1** Representative plot of small intestine lamina propria macrophages analyzed for CD88 expression together with CD4 (a), Tim-4 (b), or F4/80 (c). (d) Representative histograms of CD64 expression on the indicated populations of the monocyte-waterfall (gated as in Fig 1i), macrophage, or cDC population. Dashed line: isotype control.

**Figure S3.**
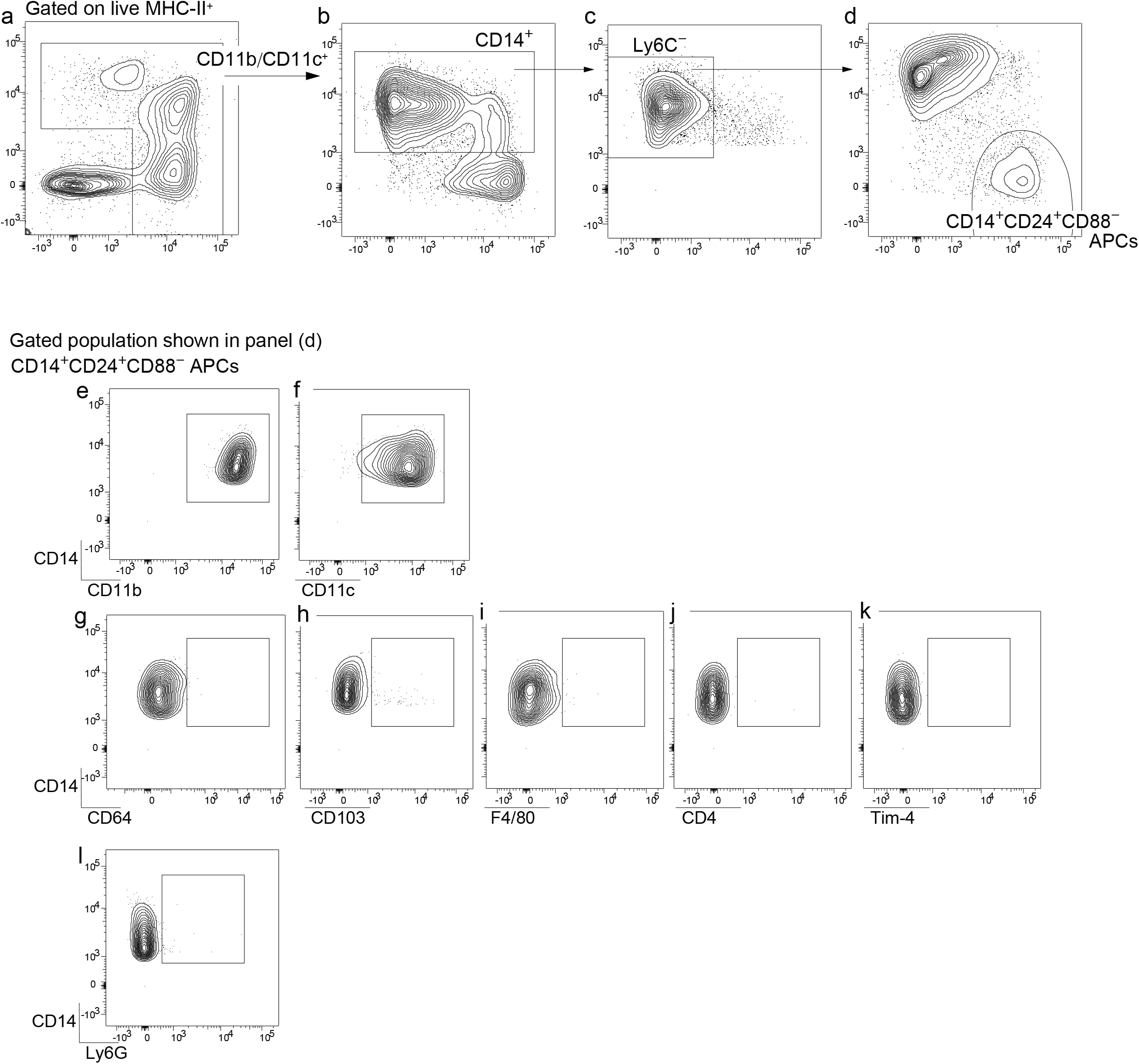
**Related to Figure 1** (a-d) Gating strategy for C-LP CD14^+^CD24^+^CD88^−^ APCs. Live MHC-II^+^ APCs as in Fig S1 were gated on populations expressing either CD11c or CD11b (a), followed by gating on CD14^+^ (b), and Ly6C^negative^ populations (c). Gated populations in (c) were subsequently analyzed for CD88 and CD24 expression to distinguish CD14^+^CD24^+^CD88^−^ APCs (d). Representative plot of C-LP CD14^+^CD24^+^CD88^−^ APCs (gated in (d)) analyzed for expression of CD11b (e), CD11c (f), CD64 (g), CD103 (h), F4/80 (i), CD4 (j), Tim-4 (k), and Ly6G (l).

**Figure S4.**
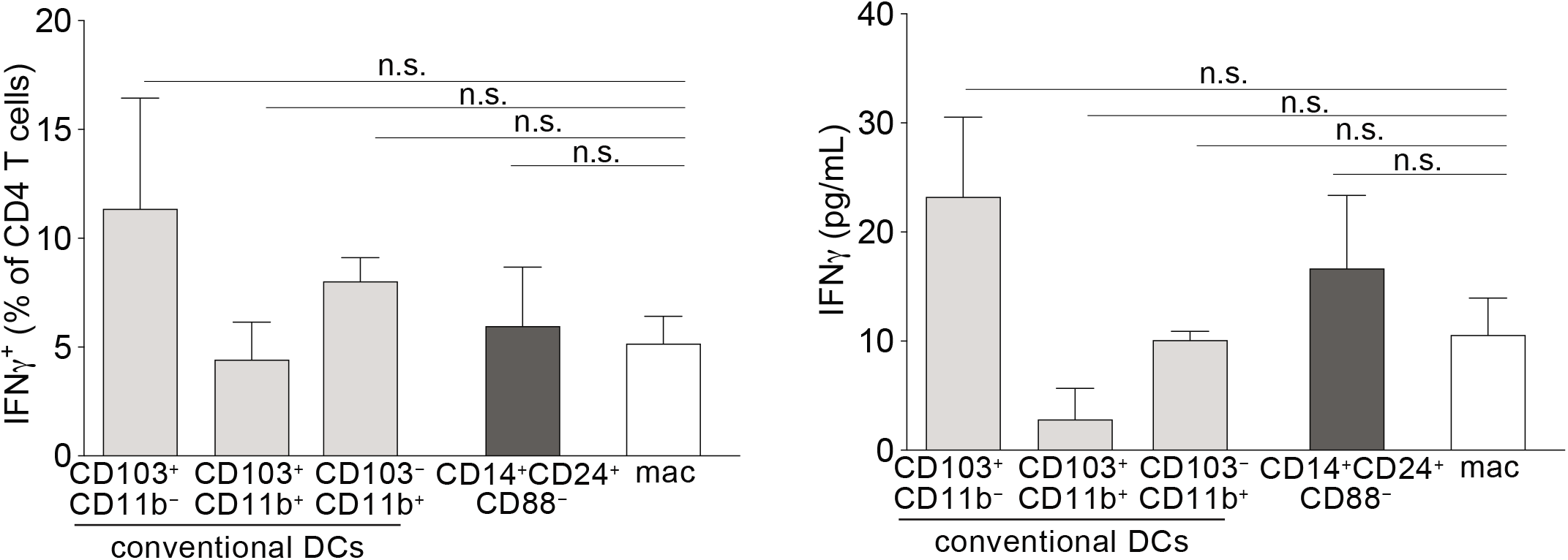
**Related to Figure 1** Percentage of IFNγ^+^ OT-II T cells (a) and IFNγ protein (b) in supernatant of co-cultures described in Fig. 1v. Error bars represent mean ± SEM. n.s. not significant (one-way ANOVA).

**Figure S5.**
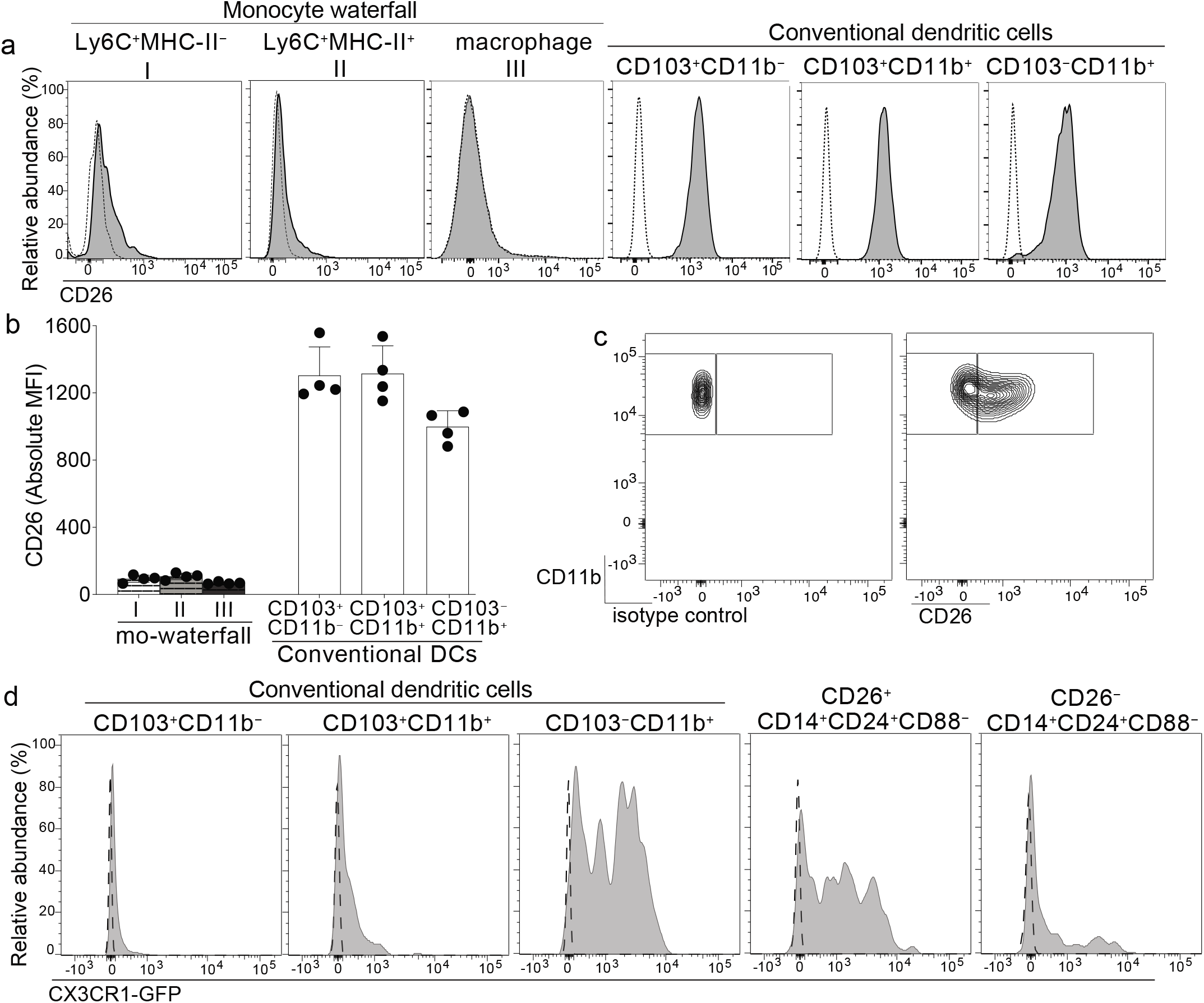
**Related to Figure 2** (a-b) CD26 expression on the indicated C-LP mononuclear phagocyte population. Representative histogram (a) and absolute MFI (b) of CD26 expression on the indicated population. Dashed line: isotype control. Each dot in (b) represents one mouse. (c) Representative plot of isotype control or CD26 expression on C-LP CD14^+^CD24^+^CD88^−^ APCs gated as in Fig 1u. (d) Representative histogram of CX3CR1-GFP expression on the indicated C-LP population. Dashed line: wild-type control.

**Figure S6.**
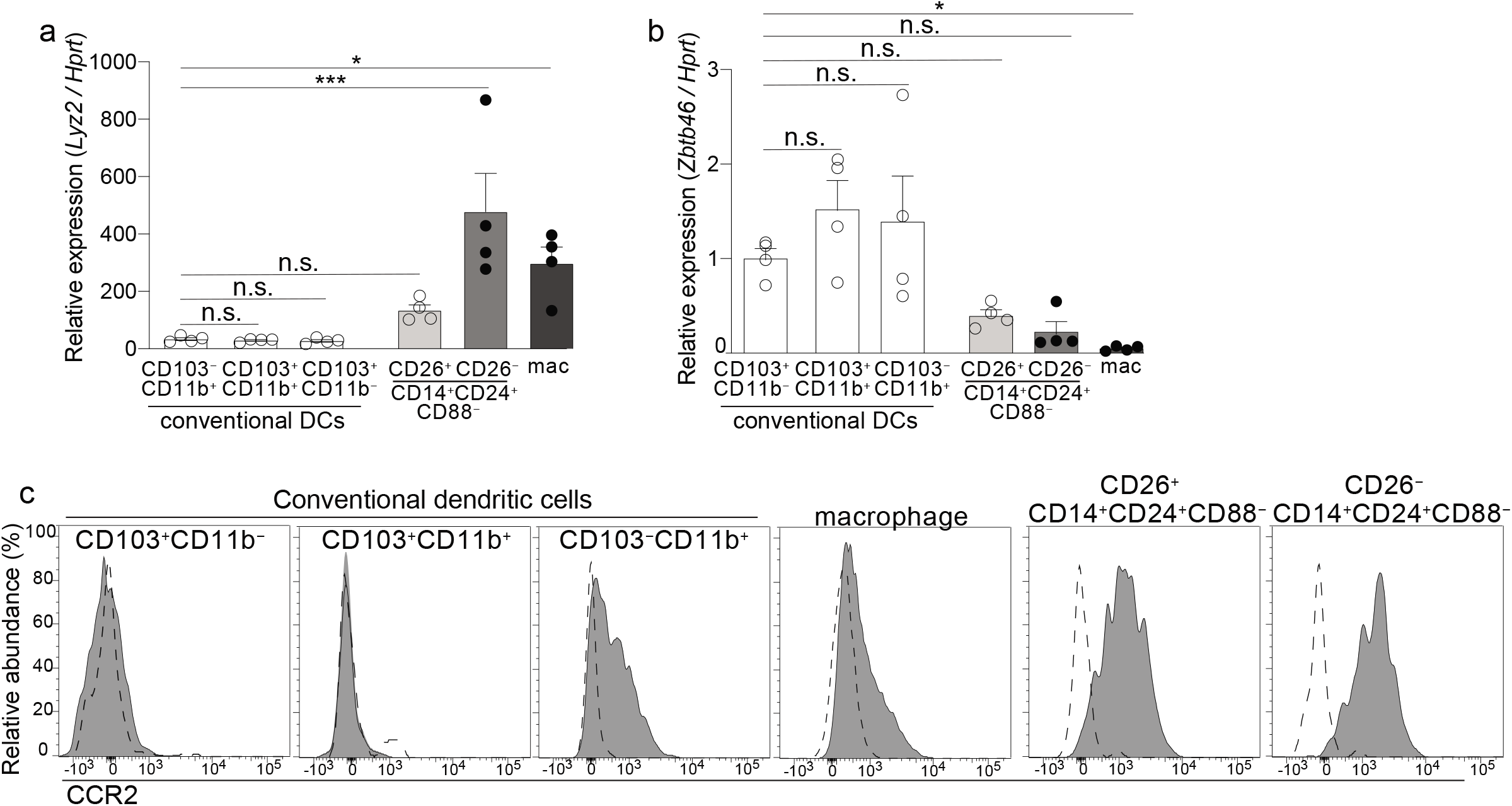
**Related to Figure 2** (a,b) Gene expression of *Lyz2* (a) or *Zbtb46* (b) expression in the indicated C-LP population. Results are relative to expression of *Hprt*. Data was combined from 3-4 independent experiments each including pooled cells from at least 5 mice for each data point. (c) Representative histogram of CCR2 expression on the indicated C-LP mononuclear phagocyte population. Error bars represent mean ± SEM. \**p* < 0.05, \*\*\**p* < 0.001, n.s. not significant (one-way ANOVA with Dunnett’s post hoc test contrasted to CD103^+^CD11b^−^ cDCs).

**Figure S7.**
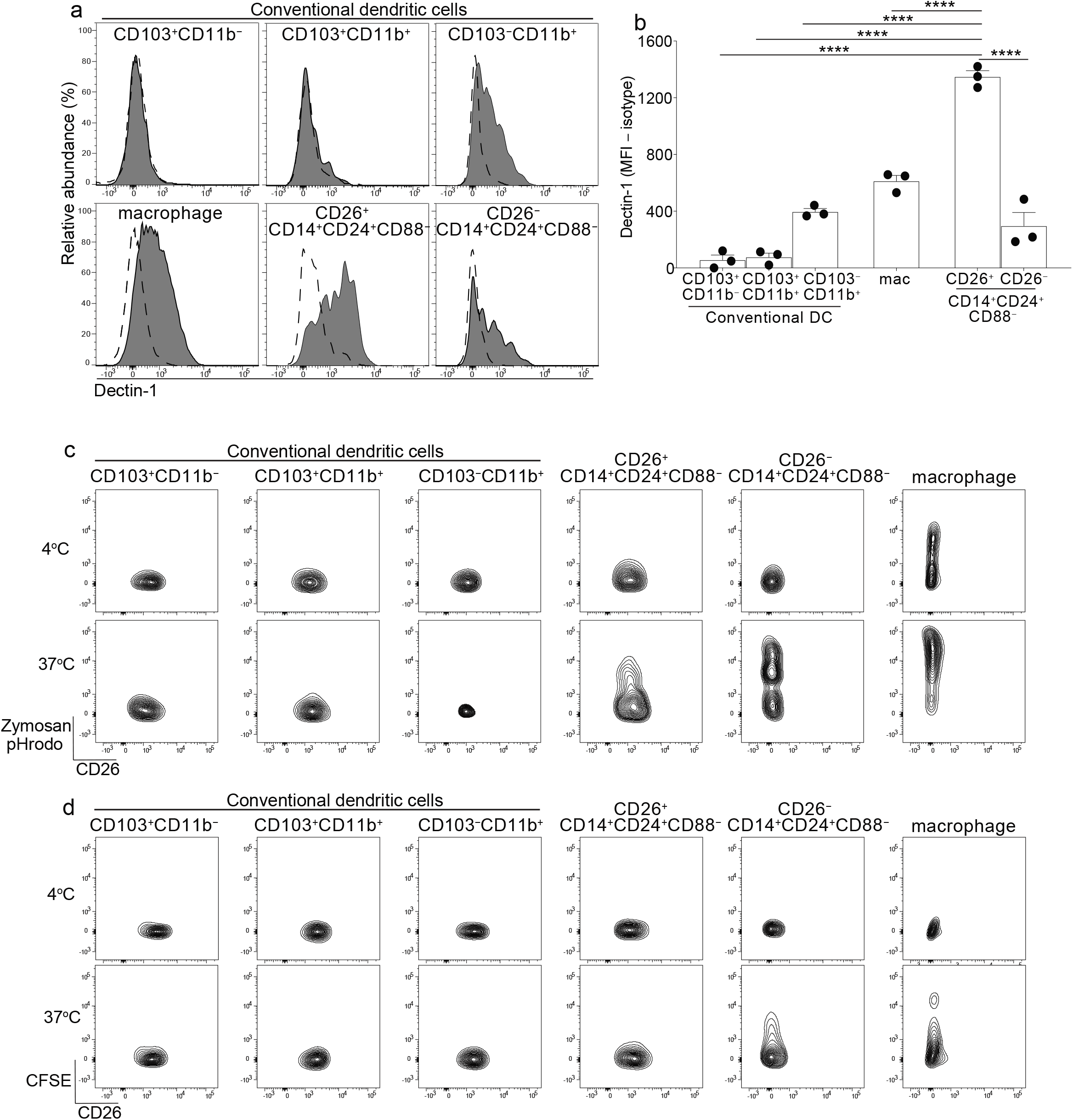
**Related to Figure 2** (a,b) Surface Dectin-1 expression on the indicated C-LP mononuclear phagocyte subset. Representative histogram (a) and MFI for all mice tested (b). Dashed line: isotype control. Each dot in (b) represents one mouse. Error bars represent mean ± SEM. (c-d) Representative plot of zymosan pHrodo bioparticle™ fluorescence or CFSE in the indicated C-LP mononuclear phagocyte subset incubated with zymosan pHrodo bioparticles™ (c) or CFSE labeled apoptotic thymocytes (d) for 1h at 4°C or 37°C. Error bars represent mean ± SEM. \*\*\*\**p* < 0.0001 (one-way ANOVA with Tukey’s post hoc test).

**Figure S8.**
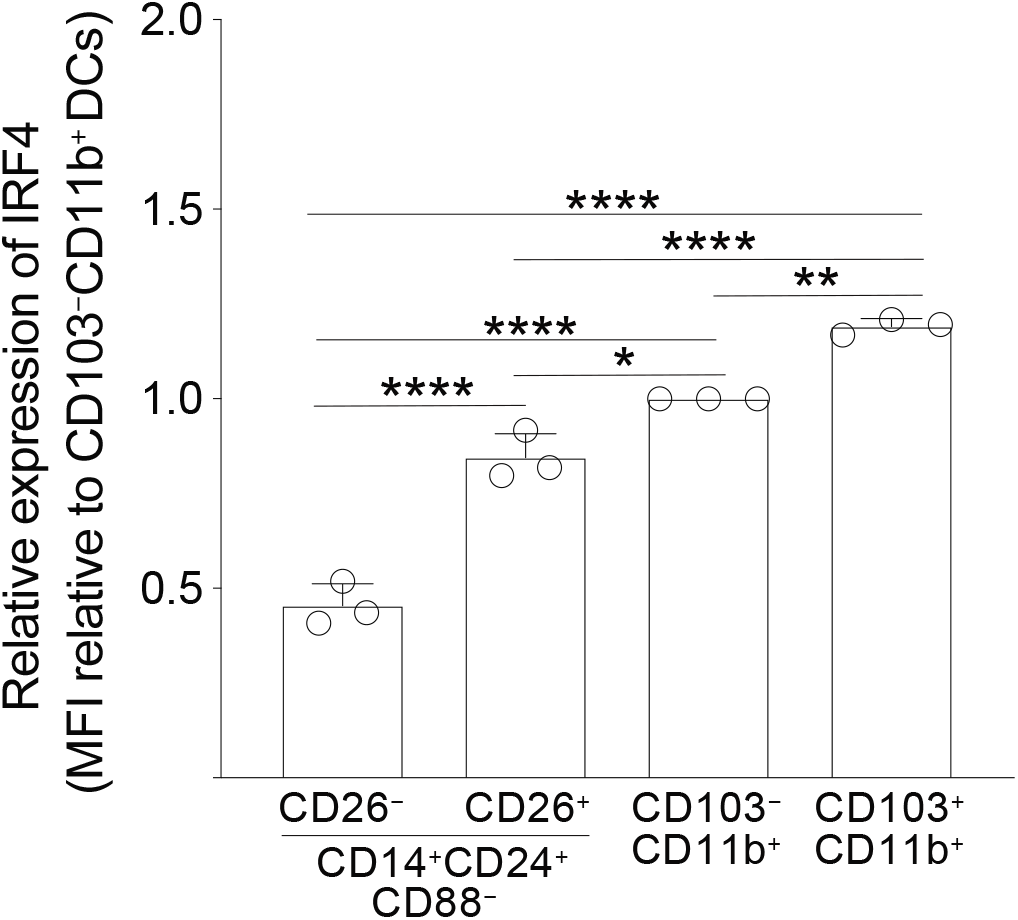
**Related to Figure 4** Relative expression (MFI) of IRF4 protein in the indicated C-LP mononuclear phagocyte subset. Each dot represents one experiment containing pooled cells from at least 2 mice. Data is combined from three independent experiments. Error bars represent mean ± SEM. \**p* < 0.05, \*\**p* < 0.01, \*\*\**p* < 0.001, \*\*\*\**p* < 0.0001 (one-way ANOVA with Tukey’s post hoc test).

**Figure S9.**
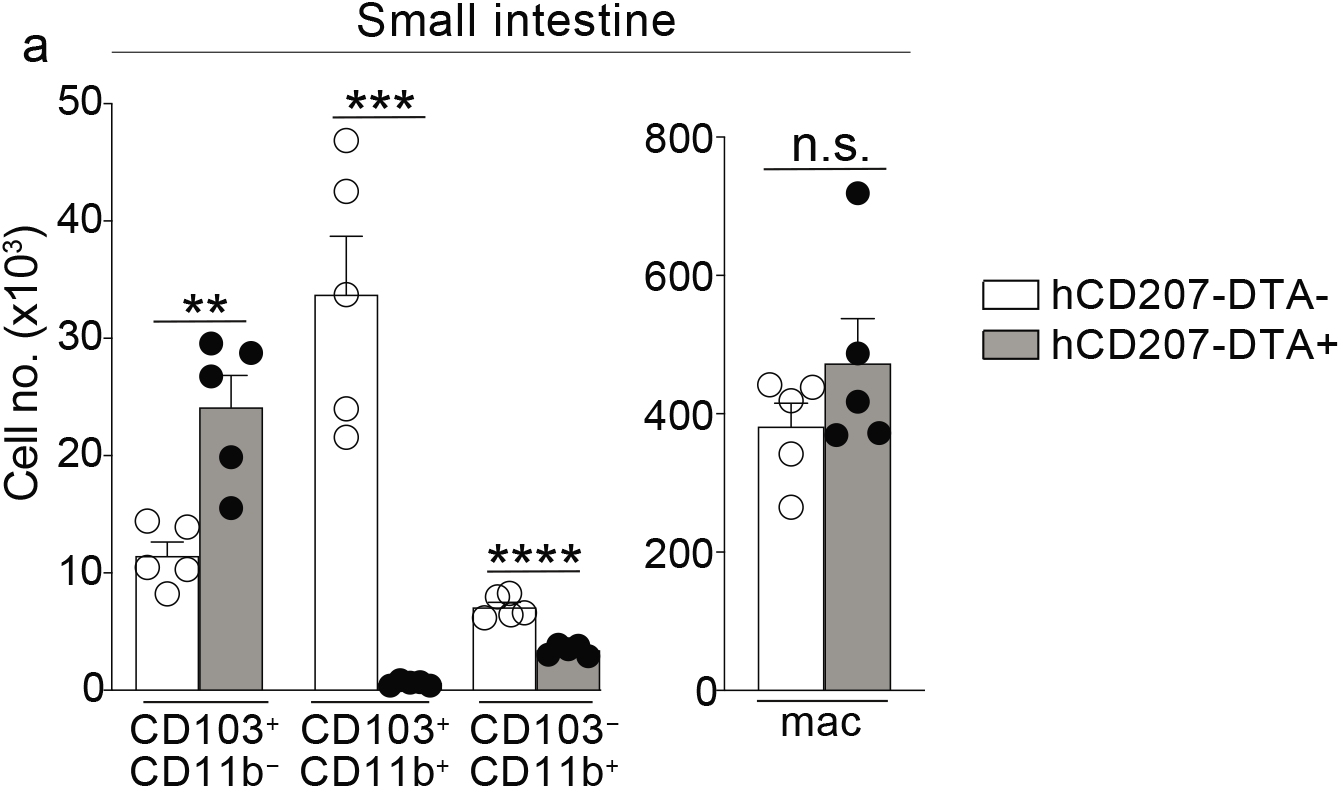
**Related to Figure 5** Absolute cell number of the indicated small intestine lamina propria mononuclear phagocyte subset from cohoused, littermate hCD207-DTA^−^ or hCD207-DTA^+^ mice. Each dot represents one mouse. Error bars represent mean ± SEM. \**p* < 0.05, \*\**p* < 0.01, \*\*\**p* < 0.001, \*\*\*\**p* < 0.0001, n.s. not significant (unpaired Student’s t test).

**Figure S10.**
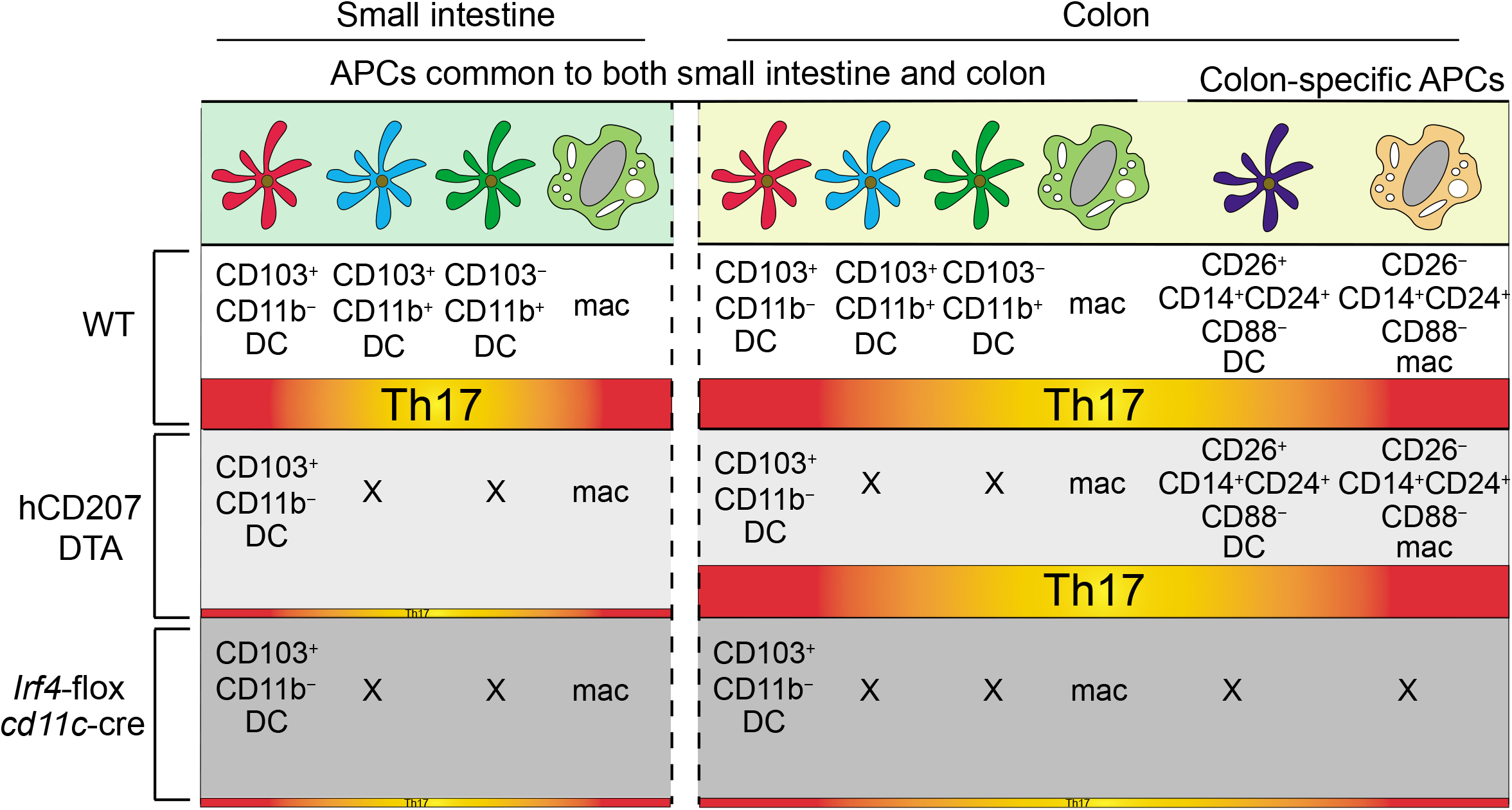
Graphical abstract. DC and macrophage subsets resident in small intestine and colon, and Th17 cells in each intestinal organ, in mice of the indicated genotype. Unlike other APC subsets, CD14^+^CD24^+^CD88^−^ APCs are specific to the colon. Colon-specific CD14^+^CD24^+^CD88^−^ APCs persist in hCD207-DTA mice, yet are ablated from *Irf4*-cko mice. This is unlike CD103^+^CD11b^+^ and CD103^−^CD11b^+^ DCs, which are depleted from both mouse strains. In the small intestine, CD103^+^CD11b^+^ and CD103^−^CD11b^+^ DCs are required for Th17 cells. This requirement is specific to small intestine, since colon-resident Th17 cells persist even in the absence of CD103^+^CD11b^+^ and CD103^−^CD11b^+^ DCs. Colon-resident Th17 cells instead require CD14^+^CD24^+^CD88^−^ APCs, which are colonspecific.

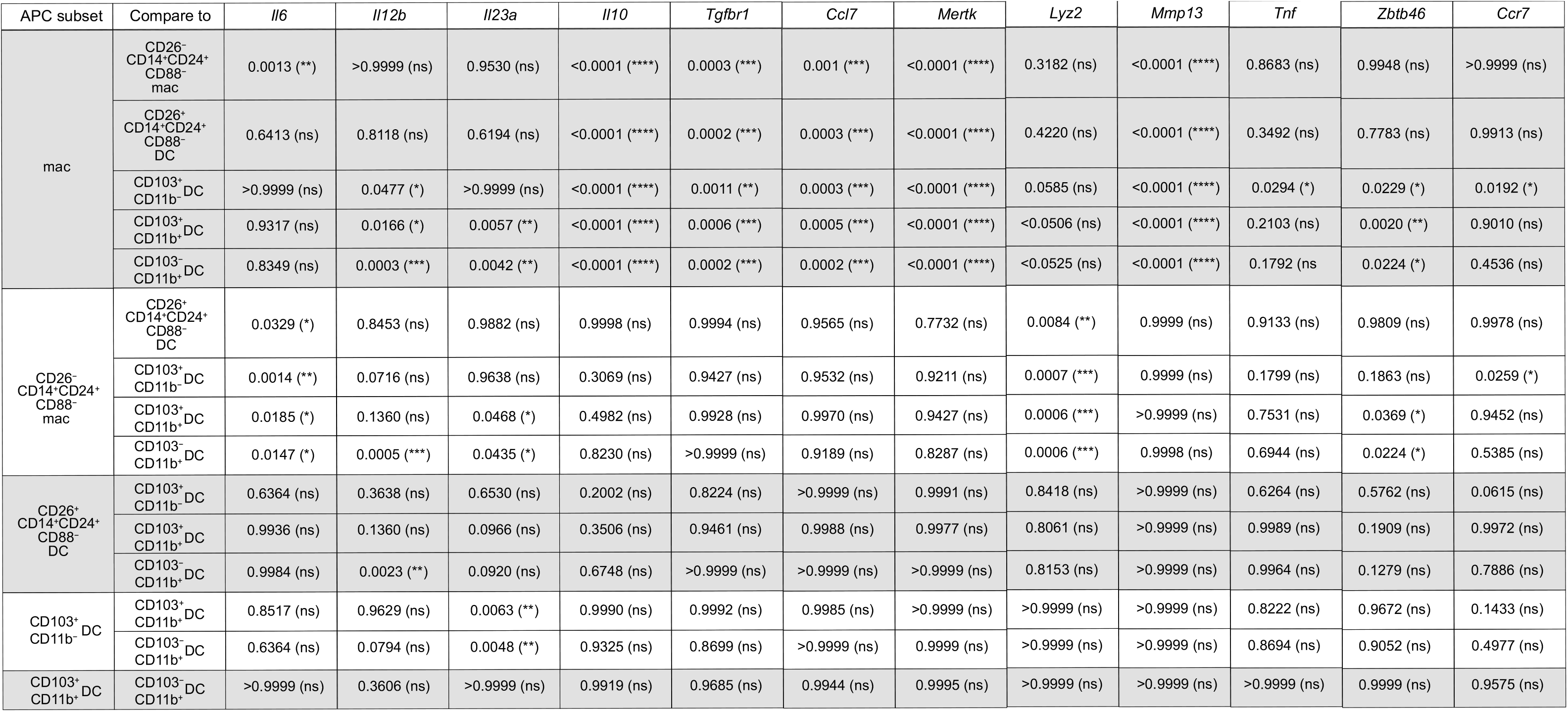
Genes Tested (Adjusted p value (significance))

